# Identification of targetable epigenetic vulnerabilities for uveal melanoma

**DOI:** 10.1101/2024.10.11.617464

**Authors:** G. Yenisehirli, S. Borges, S. Braun, A. N. Zuniga, G. I. Quintana, J. N. Kutsnetsoff, S. Rodriguez, E. V. Adis, S. Lopez, J. J. Dollar, V. Stathias, C. H. Volmar, E. Karaca, S. P. Brothers, D. C. Bilbao, J. W. Harbour, Z. M. Correa, S. Kurtenbach

**Affiliations:** Department of Ophthalmology and Sylvester Comprehensive Cancer Center, University of Miami Miller School of Medicine; Interdisciplinary Stem Cell Institute (ISCI), University of Miami Miller School of Medicine; Center for Therapeutic Innovation, University of Miami Miller School of Medicine; Department of Psychiatry and Behavioral Sciences, University of Miami Miller School of Medicine; Department of Pathology and Laboratory Medicine, University of Miami Miller School of Medicine; Department of Ophthalmology, University of Texas Southwestern Medical Center, Dallas, TX; Simmons Comprehensive Cancer Center, University of Texas Southwestern Medical Center, Dallas, TX

## Abstract

Uveal melanoma (UM) is the most common primary intraocular malignancy in adults, with a strong predilection for hepatic metastasis, occurring in approximately 50% of cases. Metastatic UM remains highly resistant to therapy and is almost invariably fatal. The strongest genetic drivers of UM metastasis are loss-of-function mutations in tumor suppressor *BAP1*, an epigenetic regulator that serves as the ubiquitin hydrolase subunit of the polycomb repressive deubiquitinase (PR-DUB) complex, and a key player in global epigenetic regulation. Inactivation of BRCA Associated Protein 1 (BAP1) has been shown to induce widespread epigenetic alterations across multiple model systems. To identify novel therapeutic strategies, we investigated whether targeting the epigenome could reveal new vulnerabilities in UM. We performed high-throughput compound screening using a curated epigenetic inhibitor library and identified BET (bromodomain and extra-terminal domain) inhibition as a particularly promising approach. Interestingly, we observed significant heterogeneity in the efficacy of different BET inhibitors in UM. While previous clinical trials with two BET inhibitors have failed to show efficacy in UM, our findings highlight substantial differences in the potency of specific BET inhibitors for this malignancy. Notably, the BET inhibitor mivebresib (ABBV-075) significantly improved survival rates by 50% in a metastatic UM xenograft mouse model and completely prevented detectable metastases in the bones, spinal cord, and brain. Unexpectedly, RNA sequencing revealed a strong transcriptional overlap between BET inhibition and histone deacetylase (HDAC) inhibition--an approach currently under clinical evaluation for UM treatment. Both BET and HDAC inhibitors reversed gene expression signatures associated with high metastatic risk and induced a neuronal differentiation-like phenotype in UM cells. Together, our findings demonstrate that UM cells exhibit a distinct vulnerability to BET inhibition and establish BET inhibitors as promising candidates for further clinical evaluation for metastatic UM.

## INTRODUCTION

Uveal melanoma (UM) is the most prevalent primary intraocular malignancy in adults, with metastases occurring in approximately half of all cases. UM metastases are highly resistant to treatment and almost uniformly lethal (1). Currently, the only FDA-approved treatment for metastatic UM is tebentafusp-tebn (Kimmtrak, Immunocore Limited), a bispecific gp100 peptide-HLA-directed CD3 T-cell engager. However, this treatment is only efficient in HLA-A*02:01-positive patients, and increases life expectancy by six months on average (2). Despite this development being a significant advancement, additional treatment strategies are urgently needed.

UM has a low mutational burden, with a mutational profile distinct from that of cutaneous and other melanomas (3). Mutually exclusive mutations in the Gq signaling pathway, most commonly in *GNAQ* or *GNA11* (4, 5), and less frequently in *PLCB4* (6) and *CYSLTR2* (7), are present in virtually all UMs (8), but also in benign ocular nevi (4, 5, 8, 9). Therefore, these mutations alone are insufficient for malignant transformation. Additional secondary mutations in either *<u>B</u>AP1* (10), *<u>S</u>F3B1* (11), or *<u>E</u>IF1AX* (12) (‘BSE’ mutations) occur in a mutually exclusive manner and are associated with high, medium, and low metastatic risk respectively (13–15). *BAP1* loss-of-function mutations are among the most significant clinical markers of high metastatic risk. Mutations in *BAP1* are typically accompanied by the loss of one copy of chromosome 3, where *BAP1* is located, resulting in the complete loss of BAP1 function (10). BAP1 is a ubiquitin carboxy-terminal hydrolase and the catalytic subunit of the polycomb repressive deubiquitinase complex (PR-DUB), which opposes PRC1 activity by removing transcriptionally repressive monoubiquitin marks from histone H2A on K119 (16–18). BAP1 depletion in various cell and animal models leads to global changes in H2AK119 ubiquitination and the epigenomic landscape (19, 20). BAP1 loss also leads to the failure of the H3K27ac histone mark accumulation at promoter sites of key lineage commitment genes, highlighting its role in the broader regulation of transcription and cell differentiation (19).

Given the role of epigenetic dysregulation in UM metastasis (21), we conducted a high-throughput screen of epigenetic modulators. We identify several new compounds with high efficacy, and highlight BET inhibition as a promising treatment angle for UM.

## RESULTS

### Epigenetic compound screening identifies new vulnerabilities in UM

Given the global epigenetic changes elicited by BAP1 loss in metastatic UM, we explored whether targeting epigenetic regulators would reveal new promising treatment angles. We performed comprehensive screening using a well-characterized compound library consisting of 932 cell-permeable, small-molecule epigenetic modulators (TargetMol, L1200), many of which are FDA-approved. We tested two *BAP1-*mutant UM cell lines (MP38 and MP46), as well as one *BAP1*-wildtype cell line (MP41) (22). The initial screen proved to be specific and identified 24 compounds that significantly reduced cell viability in at least one cell line at 1 µM after 72 h of treatment (n = 2 per compound) (fig. 1A). Most of the drug classes had low efficacy, including histone methyltransferase inhibitors (17% of compounds tested (n = 160), 0% hits), histone acetyltransferase inhibitors (7% of compounds tested (n = 68), 0% of hits), and ataxin inhibitors (18% of compounds tested (n = 167), 8% of hits (n = 2)) (fig. 1B, 1C). BET inhibitors (4% of the compound library (n = 35)), comprised 29% of the hits (n = 7), and HDAC inhibitors (7% of the compounds tested (n = 64)), were 25% of the hits (n = 6). Poly ADP Ribose Polymerase (PARP) inhibitors (n = 28) did not reduce cell viability in these cell lines (fig. 1B, supplemental fig. 1A).

**Figure 1.**
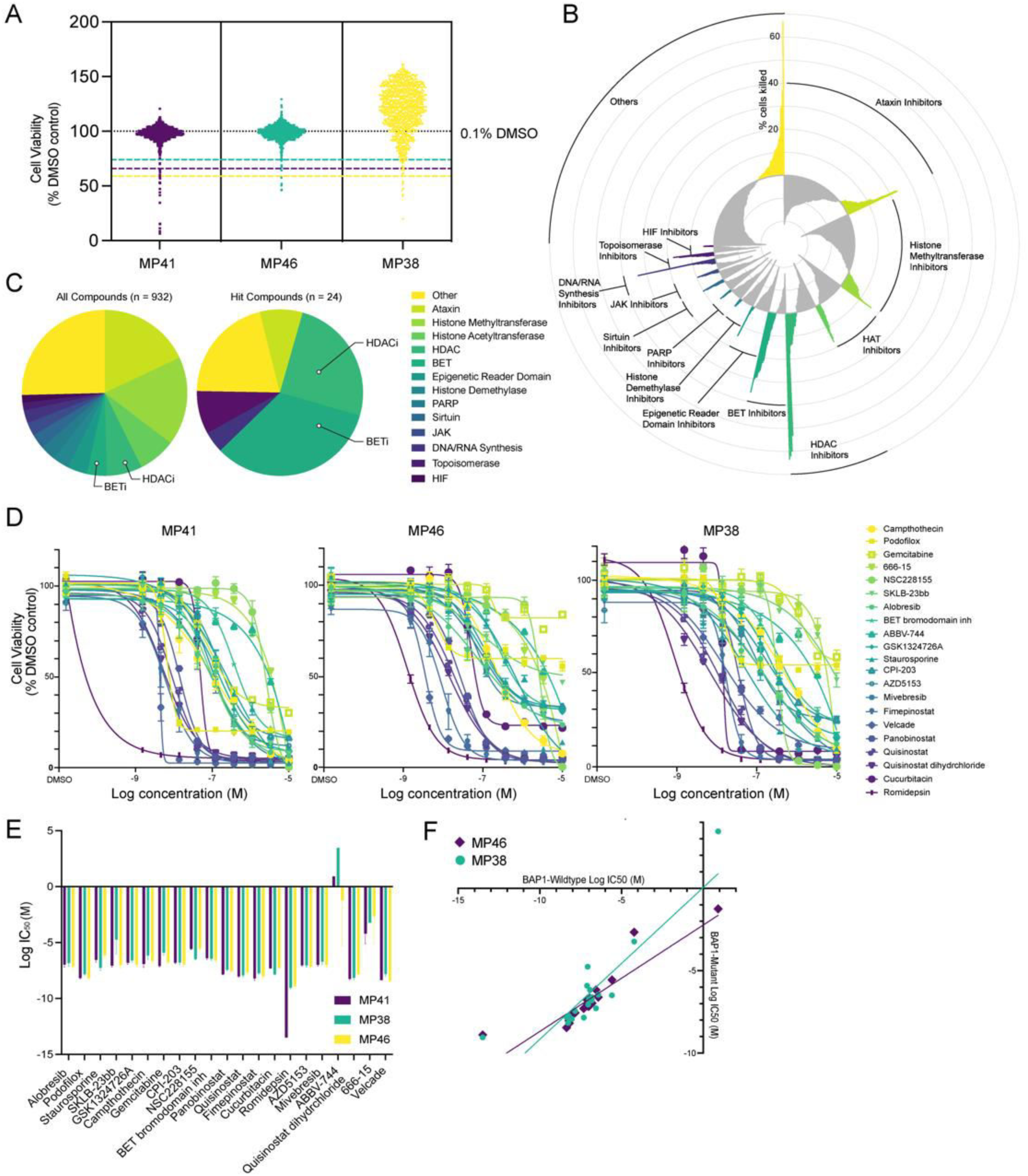
Primary screening for epigenetic compounds in UM cells. **(A)** Viability of the three UM cell lines following 72 h treatment with 932 epigenetic modulators at a concentration of 1 μM (n = 2) relative to 0.1% DMSO control. Hit cut-offs (dashed lines) were determined as the mean of the negative controls minus three standard deviations. **(B)** Radar plot showing the average difference in percentage of cell viability caused by 1 μM treatment with 932 compounds, relative to the DMSO control for the three cell lines treated with. Negative values, shown in gray, indicate ineffective compounds leading to greater cell viability than the negative control. The positive values, depicted in color, indicate compounds that induced cell death, with higher peaks indicating greater cell death. Compounds are grouped by drug mechanism of action. **(C)** Pie charts of the molecular activities of all screened compounds (n = 932) (left) and the hits identified (n = 24) (right). **(D)** Dose-response experiments for the 21 identified compounds (10 concentrations, n = 4 per concentration per cell line). Error bars represent standard deviation (SD). **(E)** LogIC_50_ values of the top hit compounds for each cell line. Error bars represent 95% confidence interval. **(F)** LogIC_50_ (M) of *BAP1* mutant cell lines (MP46 and MP38) plotted against the logIC_50_ (M) of the *BAP1* wildtype cell line (MP41) for each drug treatment.

We then tested the 21 most promising compounds in comprehensive concentration-response regimens (10 concentrations, n = 4) (fig. 1D, supplemental table 1), identifying 18 compounds with IC_50_ values less than 1 µM. The HDAC inhibitor romidepsin had the highest potency in all UM cell lines (IC_50_ ≈ 3.5 nM), even lower than that of velcade (IC_50_ ≈ 7.6 nM), a highly cytotoxic proteasome inhibitor (23) used as a positive control in this screen.

Of the 18 compounds with an IC_50_ of less than 1 µM, 13 were HDAC and BET inhibitors and five compounds targeted other mechanisms. Of the latter, gemcitabine (IC_50_ ≈ 493 nM) and staurosporine (IC_50_ ≈ 336 nM), have previously been shown to induce apoptosis in UM cells (24, 25). Camptothecin (IC_50_ ≈ 334 nM) (topoisomerase I inhibitor (26)), podofilox (IC_50_ ≈ 9.36 nM) (microtubule destabilizer (27)) and cucurbitacin B (IC_50_ ≈ 37.9 nM) (inhibitor of AKT, HIF1a, and STAT3 (28)), to our knowledge, have not previously been tested for UM. All compounds had similar IC_50_ values in the cell lines tested, despite their genetic differences, namely MP41 being *BAP1*-wildtype and MP38 and MP46 being *BAP1*-mutant (fig. 1E, 1F). We further tested for synergy between romidepsin and quisinostat with the other 16 compounds. However, despite these compounds targeting diverse epigenetic pathways, we did not find significant synergy (supplemental fig. 2).

### HDAC inhibition in uveal melanoma cells

HDAC inhibition has been explored in numerous studies and clinical trials, so far with limited success (29–33). There are 11 HDAC isoforms in humans that function in numerous protein complexes and have diverse biological functions, and it is unclear which HDACs are the most promising to specifically target in UM (34, 35). Romidepsin demonstrated the greatest potency, suggesting that inhibition of class I HDACs may be a vulnerability for UM, as romidepsin specifically inhibits class I HDACs (HDAC1, 2, 3, and 8) (fig. 2B). Although no specific inhibitors for HDAC1 and HDAC2 exist, we tested the HDAC3 inhibitor RGFP966 (TargetMol, T1762) and the HDAC8 inhibitor PCI-34051 (TargetMol, T6325) and found that neither was potent, neither alone or in combination (supplemental fig. 1B). We tested romidepsin from two different sources (TargetMol T6006, Sigma SML1175) and included an additional primary *BAP1*-mutant UM cell line we generated (UMM66) (fig. 2A). Both romidepsin batches showed similar potency in all UM cell lines, including UMM66 cells (IC_50_ = 2.4 - 5.7 nM). Together, these data highlight romidepsin as the most potent compound in this *in vitro* screen, and specific inhibition of class I HDACs, likely acting through HDAC1 and HDAC2, as a potential vulnerability of UM.

**Figure 2.**
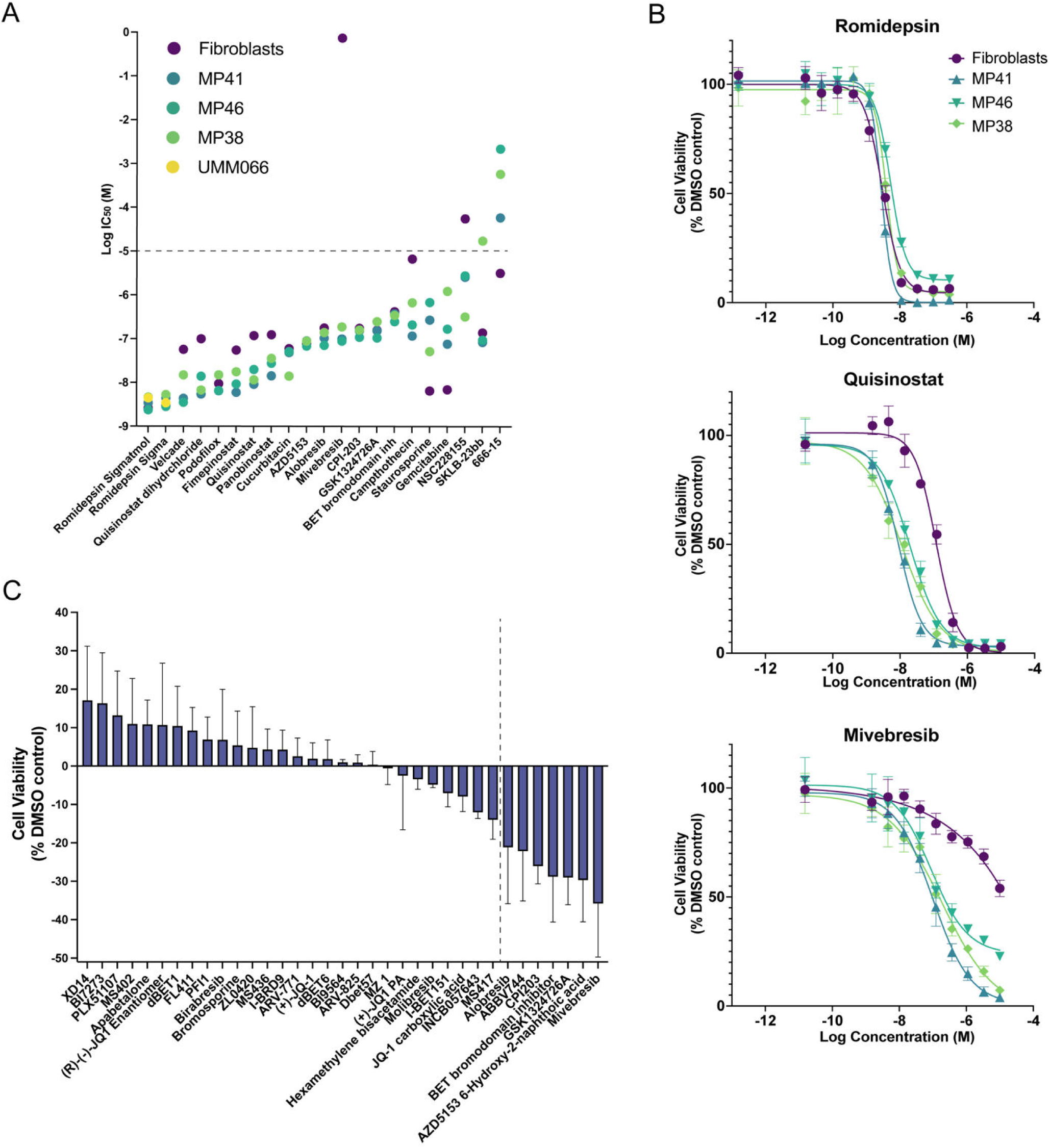
IC_50_ and dose-response curves of lead compounds in UM cell lines and normal fibroblasts. **(A)** LogIC_50_ values of the lead compounds tested in UM cell lines and normal fibroblasts. The dotted line indicates the highest concentration of drug tested (10 μM); hence, for the values above, the IC_50_ is likely not calculated accurately. n = 4 replicates for each concentration were tested. **(B)** Concentration-response curves of the top candidates (romidepsin, mivebresib, and quisinostat) for UM cell lines and WS1 fibroblasts. n = 4 per concentrations tested. Error bars indicate SD. **(C)** Average difference in percent cell viability of three UM cell lines (MP41, MP38, MP46) relative to the DMSO control of all BET inhibitor drugs (1 μM) tested in primary screen. Error bars represent standard error of mean. Compounds to the right of the dotted line were identified as hits inducing significant cell death in the primary screen.

### BET inhibition in uveal melanoma cells

To explore non-specific toxicities, we performed viability assays on a non-cancerous WS1 fibroblast cell line. Fimepinostat (fibroblast IC_50_ ≈ 55 nM, UM IC_50_ ≈ 11 nM) and panobinostat (fibroblast IC_50_ ≈ 124 nM, UM IC_50_ ≈ 26 nM) demonstrated 4- to 5-fold lower toxicity to non-transformed cells. Quisinostat had an approximately 9 times higher IC_50_ for non-cancerous cells (fibroblast IC_50_ ≈ 118 nM) than for UM cells (UM IC_50_ ≈ 14 nM) (fig. 2A). Other drugs with lower cytotoxicity to normal cells included velcade (fibroblast IC_50_ ≈ 57 nM, UM IC_50_ ≈ 8 nM) and campthothecin (fibroblast IC_50_ ≈ 7 µM, UM IC_50_ ≈ 334 nM).

Most of the 35 BET inhibitors were not efficient in reducing UM cell viability at the 1 μM concentration tested in the primary screen (fig. 2C). However, the BET inhibitor mivebresib showed minimal toxicity to normal fibroblasts (IC_50_ > 10 μM), while being among the most potent BET inhibitors tested (IC_50_ ≈ 125 nM). (fig. 2B, 2C). This data highlights a significant heterogeneity in the responses of UM cells to different BET inhibitors.

Although treatment of the primary tumor UM has a high rate of success, approximately half of all patients develop fatal metastases, primarily in the liver. Therefore, we tested our lead compounds in a mouse model to assess their ability to reduce metastatic growth. We included mivebresib and HDAC inhibitor quisinostat due to their potency on UM cells and lower toxicity to normal fibroblasts, as well as the HDAC inhibitor romidepsin, which is FDA approved. We tested various UM cell lines and found that MP41 cells metastasize predominantly to the liver when injected into the tail vein. MP41 is *BAP1*-wildtype, and was derived from an aggressive UM case that had spread to multiple organs and has features of *BAP1*-mutant UM, including monosomy 3 (36). We deemed this model as most suitable to explore the inhibition of metastatic growth in the liver, as we did not find significant differences between MP41 and the *BAP1*-mutant cell lines MP46 and MP38 regarding drug sensitivity,

Toxicity assays were conducted initially to determine optimal drug doses. We labeled MP41 cells with luciferase for *in vivo* and *ex vivo* monitoring of organs for metastatic disease. Seven days after cell injection, drug treatments were initiated to determine the efficacy of each treatment in slowing metastatic growth (fig. 3A). Quisinostat and romidepsin treatments did not significantly improve survival rates in comparison with the vehicle group (p > 0.10), with median survival rates between 83-89 days after tumor cell injection (fig. 3B). Mivebresib treatment increased median survival by 50%, to 121 days (p = 0.01) (fig. 3B). *Ex vivo* IVIS imaging revealed that mivebresib prevented metastases to the femur and spinal cord, which were detected in all other experimental groups (fig. 3C, 3D).

**Figure 3.**
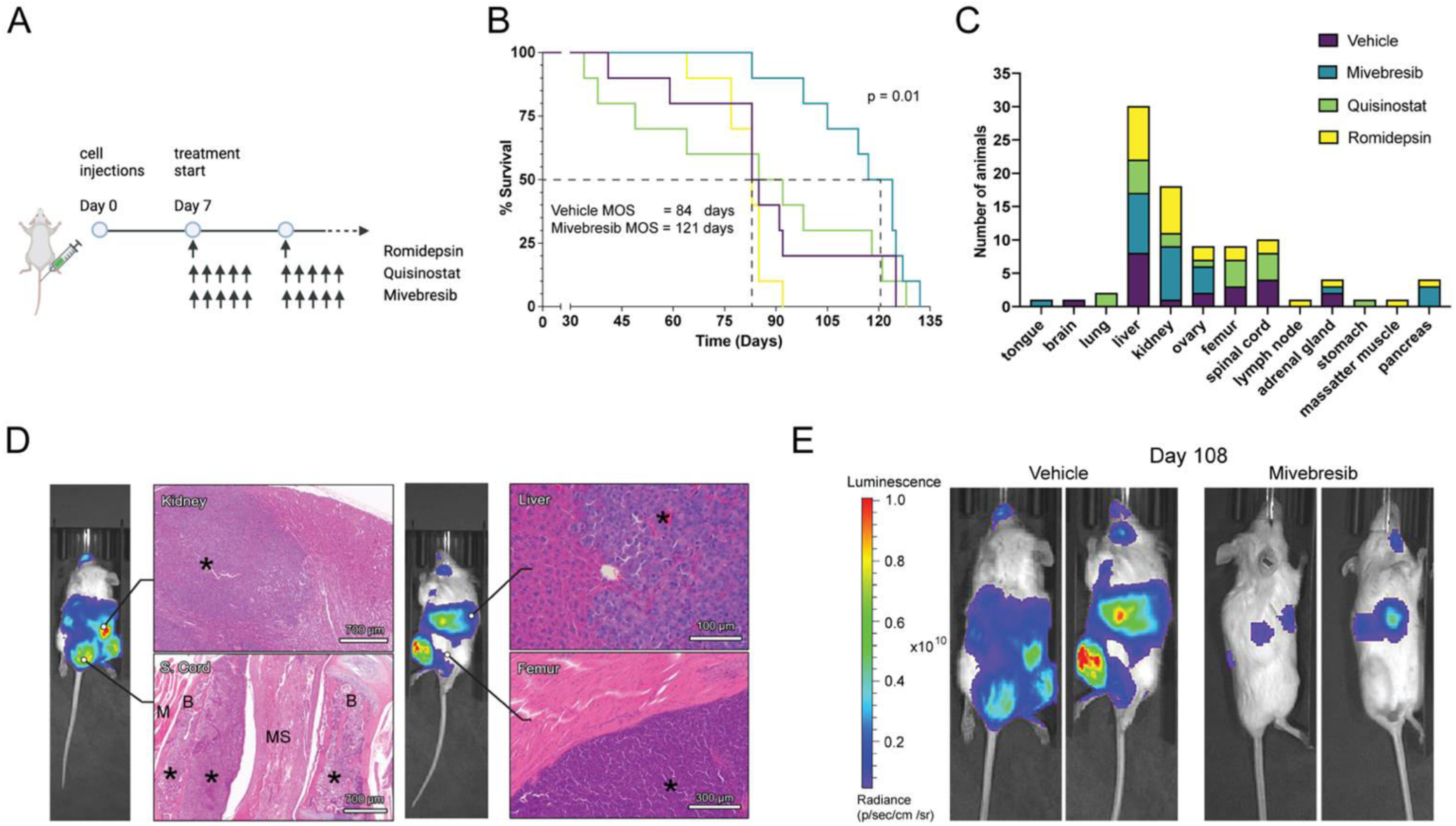
BET inhibition reduces metastatic UM growth *in vivo*. **(A)** Experimental outline and timeline of treatments in the metastatic UM mouse model. **(B)** Percent survival of mice in each treatment group (n = 10 per group) over the course of 140 days. **(C)** Bar graph depicting the number of mice in each treatment group with metastatic foci detected in different organs. **(D)** Representative histopathological images of kidney, spinal cord (S. cord), liver, and femur metastases from the vehicle-treated group. (* = tumor cells; M = muscle; B = bone; MS = medulla spinalis). **(E)** Representative IVIS images of the vehicle and mivebresib treatment groups on day 108. Luminescence/radiance in p/sec/cm^2^/sr.

To test whether long-term *in vivo* treatments led to the development of drug-resistant metastases, we extracted UM cells from mice liver metastases from all treatment groups and performed concentration-response testing with romidepsin, quisinostat, and mivebresib. No significant resistance was detected in any of the treatment groups relative to cells extracted from mice in the vehicle treatment group (supplemental fig. 3).

### Transcriptomic changes associated with HDAC and BET inhibition

To elucidate the mechanistic differences of HDAC and BET inhibition in UM, we performed bulk RNA sequencing on MP41 (*BAP1* wildtype) and MP46 (*BAP1* mutant) cell lines after 24 hours of treatment with drug concentrations that resulted in morphological changes without excessive cell death. Romidepsin, quisinostat, and mivebresib each induced unique morphological changes in MP41 cells, with both HDAC inhibitors causing a flattened morphology, whereas mivebresib-treated cells displayed mixed morphologies including flat and spindle-shaped cells (fig. 4A). Similar changes were observed in both cell lines, with unique gene expression changes for each compound and a clear separation by principle component analysis (PCA) (fig. 4B, 4C; supplemental fig. 4B, 4C). Both HDAC inhibitors resulted in an overall increase in gene expression (fig. 4D; supplemental fig. 4D), which is consistent with HDAC inhibitors leading to increased histone acetylation and chromatin accessibility (37). Most of the gene expression changes in quininostat-treated cells overlapped with those observed in romidepsin-treated cells. However, although romidepsin primarily inhibits class I HDACs, it caused more gene expression changes at the same concentration than quisinostat (fig. 4D, 4E; supplemental fig. 4D, 4E). Mivebresib treatment, on the other hand, resulted in more downregulated than upregulated genes in both cell lines (fig. 4D, 4E, supplemental fig. 4D, 4E), in concordance with BET inhibitors preventing the binding of bromodomain (BRD) proteins to acetylated histones, which typically initiate transcription by recruiting transcriptional machinery to acetylated sites (38, 39).

**Figure 4.**
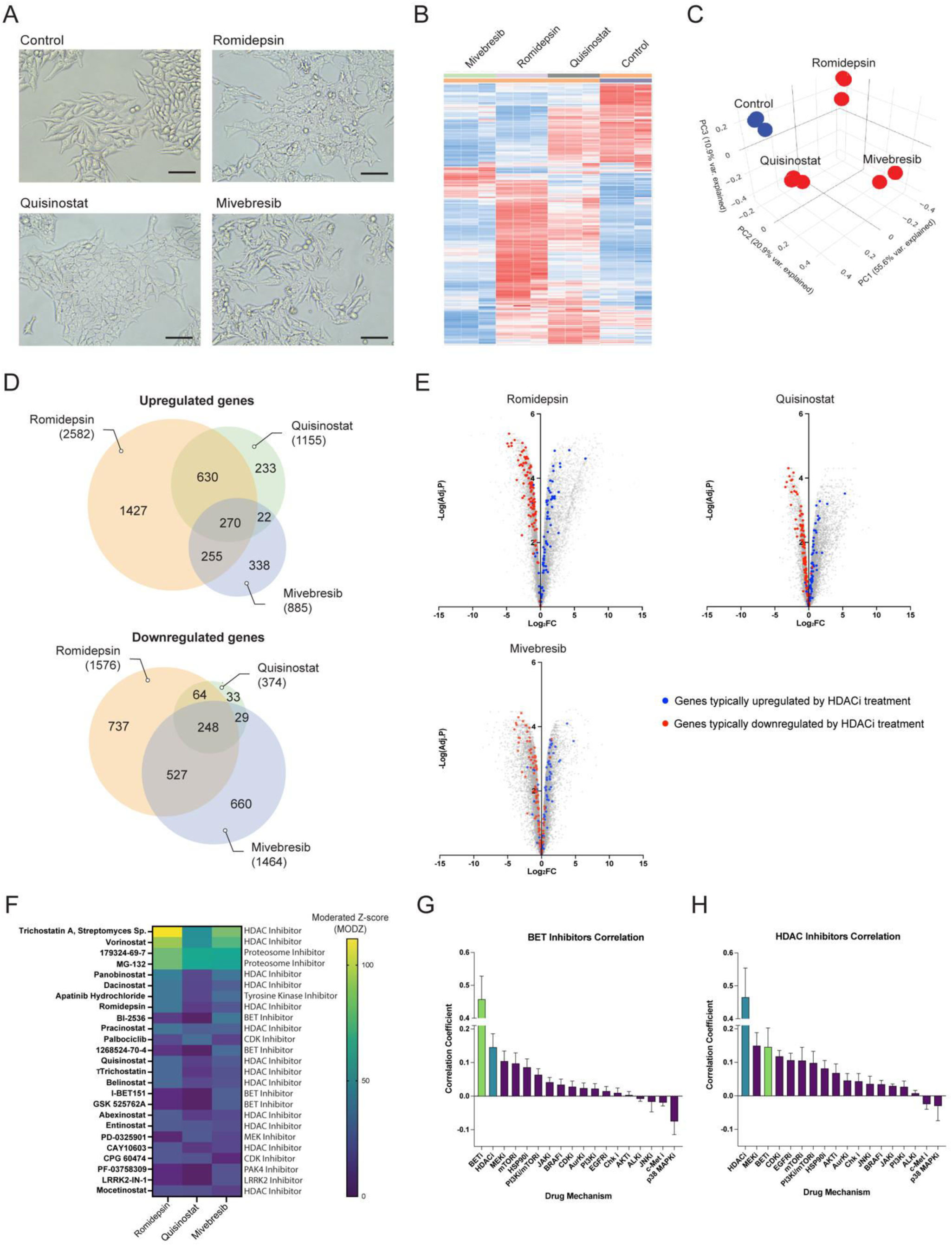
Gene expression changes following BET and HDAC inhibition. **(A)** Images of MP41 cells treated with each compound for 24 hours. Scale bar = 100 µm. **(B)** Heatmap clustering of changes in gene expression per treatment group (n = 3 per condition). **(C)** PCA analysis of replicates for each treatment in MP41 cells. **(D)** Venn diagram depicting overlaps between the treatment groups of significantly upregulated and downregulated genes in MP41 cells (Adj. P < 0.05, | log_2_ FC | > 1.5). **(E)** Volcano plot of changes in gene expression in MP41 cells relative to the control for each treatment group. Blue and red dots are the 180 genes found to be consistently dysregulated as a result of eight HDAC inhibitor treatments in iLINCS. Blue dots are genes that were consistently upregulated by HDAC inhibitor treatment (n = 77), while red dots are genes that were consistently downregulated (n = 103). **(F)** Heatmap of moderated Z-score (MODZ) of compounds inducing similar gene expression signatures to MP41 cells treated with romidepsin, quisinostat, and mivebresib using iLINCS connected perturbation analysis. Higher MODZ indicates greater similarity. **(G)** Bar graph of the average correlation coefficient of drugs with each mechanism to HDAC inhibitors. Error bars represent SEM. **(H)** Bar graph of the average correlation coefficient of drugs with each mechanism to BET inhibitors. Error bars represent SEM.

Despite their different mechanisms, we found a significant overlap in gene expression changes elicited by HDAC and BET inhibitors (fig. 4D, supplemental fig. 4D). To further investigate this finding, we compiled a list of genes consistently up- and down-regulated by HDAC inhibitors across various cancers using the Library of Integrated Network-based Cellular Signature (iLINCS) (40) database, and found that most of these genes were not only up- and down-regulated by HDAC inhibitor treatments in UM cells, but also following BET inhibition with mivebresib (fig. 4E, supplemental fig. 4E). We performed an iLINCS connected perturbations analysis, which included gene signatures from various cancer and cell models, and found that mivebresib treatment of UM cells causes a gene expression shift that is most similar to HDAC inhibitors (fig. 4E, 4F; supplemental fig. 4E, 4F). Similary, global analysis of similarities between compound classes revealed BET inhibition to be most similar to HDAC inhibition (*r* = 0.1458) compared to other classes (fig. 4G). HDAC inhibition was also most similar to MEK (*r* = 0.1494) and BET inhibition (fig. 4H).

Together, these data show that while BET inhibition may be less toxic and more efficient at reducing growth of metastatic UM, the gene expression changes elicited by BET and HDAC inhibitors have a significant overlap.

### HDAC and BET inhibition reverse transcriptomic signatures associated with high metastatic risk

Clinically, UM can be accurately stratified into metastatic risk groups, namely class 1 (low-risk) and class 2 (high-risk), using a gene expression panel of 12 genes (41–43). An additional biomarker of high metastatic risk for both class 1 and class 2 UM is the expression of *PRAME* (44–46). While MP41 cells are *BAP1* wildtype, we found that they express many class 2 markers at similar levels as the *BAP1* mutant cell line MP46. We found that treatment of MP41 and MP46 UM cells with HDAC and BET inhibitors reversed class 2 signature genes, with high-risk biomarkers such as *HTR2B* and *PRAME* being downregulated (fig. 5A, 5B). Accordingly, genes with low expression in class 2 tumors, such as *ROBO1* and *LMCD1,* were upregulated following treatment in both cell lines (fig. 5A, 5B).

**Figure 5.**
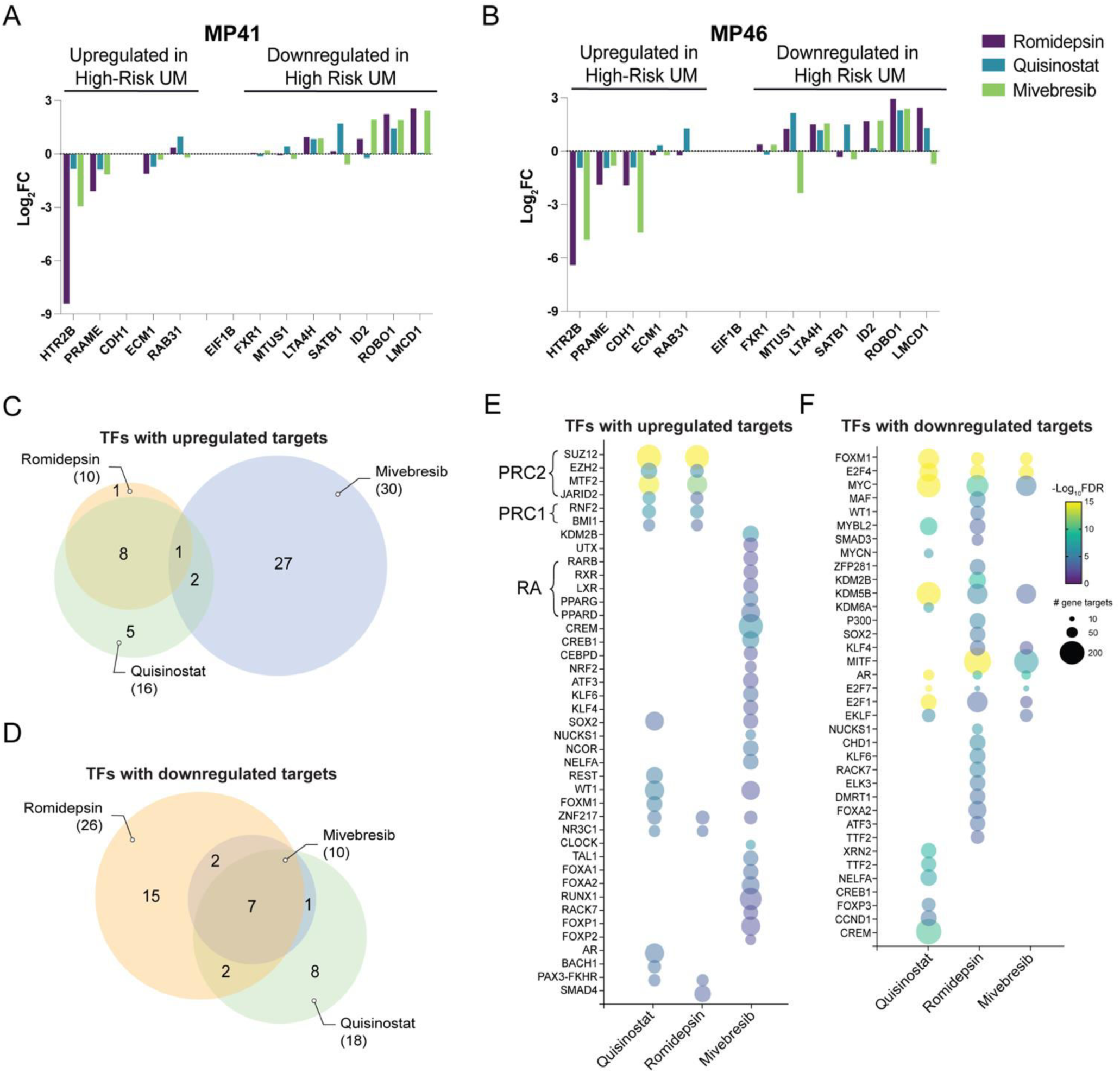
BET and HDAC inhibition act through unique mechanisms with overlapping pathway changes. **(A)** Changes in gene expression (log_2_ FC) of genes associated with high-risk UM in drug-treated in MP41 and **(B)** MP46 cells. **(C)** Venn diagram showing overlaps in predicted transcription factors with upregulated and **(D)** downregulated gene targets, inferred by gene expression changes induced by each treatment in MP41 cells. **(E)** Bubble plot of the top predicted transcription factors with upregulated and **(F)** downregulated gene targets for each treatment in MP41 cells. Color scheme indicates -Log_10_FDR of each predicted transcription factor, and bubble size is determined by the number of corresponding gene targets.

ChIP Enrichment Analysis (ChEA) (47) showed that in both MP46 and MP41 cells, the most prominent increase in gene expression following HDAC treatments were targets of the polycomb repressive complex (PRC) 1 (RNF2, BMI1) and PRC2 (SUZ12, EZH2, and cofactors MTF2, JARID2), indicating a loss of PRC activity (fig. 5E, 6J; supplemental fig. 5C). In MP41 cells, the top differential transcription factor activity for all treatments was FOXM1, whose target genes were significantly downregulated in all treatment groups (fig. 5F). FOXM1 activity is associated with a more aggressive UM phenotype, and silencing FOXM1 suppresses UM proliferation, migration, and invasion (48). Other transcription factors whose targets were downregulated in all groups included E2F family members, MYC, and the histone demethylase KDM5B (fig. 5F, supplemental fig. 5D). Although there was significant overlap in the transcription factors whose target genes were downregulated in all treatment groups, we found a large group of unique transcription factors whose activities were upregulated by mivebresib treatment (fig. 5C, 5D; supplemental fig. 5A, 5B). These factors include retinoic acid receptors RXR and RARβ and their binding partners LXR, PPARγ, and PPARδ (fig. 5E), which regulate pathways involved in neuronal differentiation (49–51). Additionally, the mivebresib treatment group exhibited unique stress-related signaling via NRF2, KLF6, and ATF3 (fig. 5E, supplemental fig. 5C).

### BET and HDAC inhibition induce a neuronal phenotype in UM cells

We observed that several genes associated with a neuronal cell identity, including *NEFM* (Neuronal Filament Medium), *SYN1* (Synapsin 1), and *NGFR* (Nerve Growth Factor Receptor), were upregulated in both MP41 and MP46 cells following HDAC or BET inhibitor treatment (fig. 6A, 6B). Neural crest and melanocytic identity genes, including *SOX10*, *MLANA*, and *MITF,* were highly downregulated (fig. 6A, 6B). Upregulation of neuronal genes in drug-treated MP41 cells was verified by immunoblotting and immunofluorescence staining of *β*-tubulin III (TUBB3) and Synapsin 1 (SYN1) (fig. 6C-F, supplemental material A, B).

**Figure 6.**
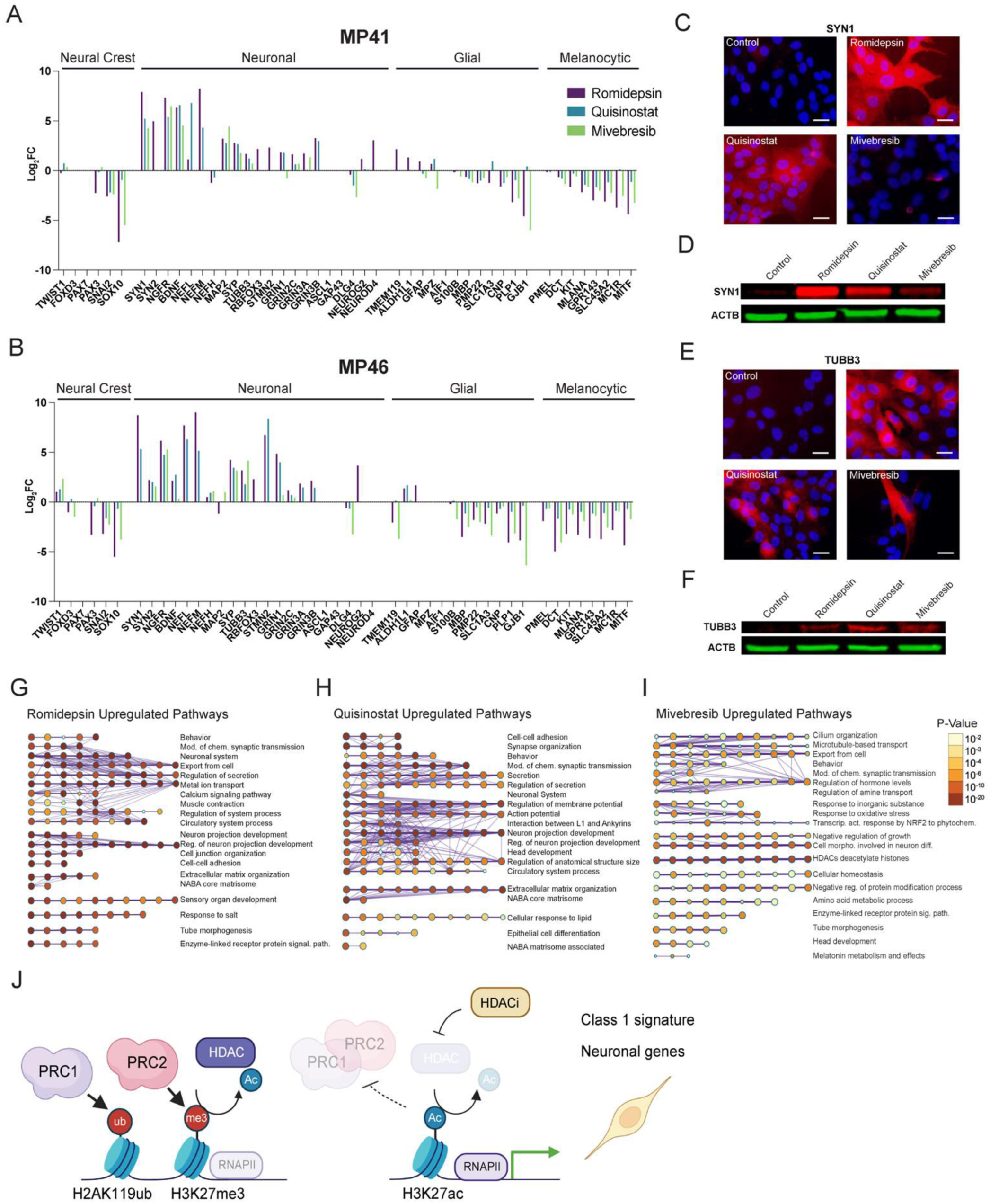
BET and HDAC inhibition induce a neuronal phenotype in UM cells. **(A)** Changes in the expression (log2 FC) of genes associated with some neural-crest-derived cell identities in drug-treated MP41 and (B) MP46 cells. (C) Immunofluorescence images of 24-hour drug treated MP41 cells. Red fluorescence is SYN1 and blue is DAPI. Scale bar = 25 μM. (D) Immunoblot of SYN1 in 24-hour drug treated cells with a *β*-Actin (ACTB) control. (E) Immunofluorescence images of 24-hour drug treated MP41 cells. Red fluorescence is TUBB3 and blue is DAPI. Scale bar = 25 μM. (F) Immunoblot of TUBB3 in 24-hour drug treated cells with a *β*-Actin (ACTB) control. (G-I) Gene interaction networks of upregulated pathways in MP41 cells predicted from significantly upregulated genes (log2 FC > 1.5, adj. p < 0.05) in each treatment group show enrichment for neuronal pathways. (J) Schemetic representation of HDAC inhibition impairing PRC activity, leading to elevated expression of PRC target genes, including neuronal genes and those associated with a class 1 phenotype.

Pathway analysis revealed the upregulation of several neuronal pathways following treatments, including synaptic transmission, neuronal projection, action potential, and neuronal differentiation (fig. 6G-I; supplemental fig. 5E-G). Compared to HDAC inhibitors, BET inhibition activated additional pathways involved in the stress response, including NRF2 signaling (fig. 6I; supplemental fig. 5G). All drug treatments induced downregulation of pathways involving DNA replication, cell growth, and proliferation (supplemental fig. 6).

Together, these data indicate that HDAC and BET inhibition induce a phenotypic identity switch, pushing cells towards a lower metastatic risk gene expression signature and neuronal cell identity (fig. 6J).

## DISCUSSION

Treatment options for metastatic UM are limited, with the only FDA-approved drug prolonging overall survival by only six months on average for a subset of patients. Here, we utilized an epigenetic compound screen to identify new vulnerabilities that target the epigenome of UM, as most metastatic UM’s harbor mutations in the epigenetic regulator *BAP1*, accompanied by global epigenetic changes. We show that HDAC and BET inhibitors were the most efficacious compound classes *in vitro*. We previously showed that PARP inhibition can reduce the metastatic spread in a mouse model of UM (44). However, our *in vitro* experiments did not identify PARP inhibitors as a potent drug class (fig. 1B, 1C; supplemental fig. 1A), indicating that PARP inhibition acts through mechanisms other than reducing cell viability *in vivo*.

HDAC inhibitors are widely considered for treatment of UM (30, 33), however, with limited clinical success so far. The class I HDAC inhibitor romidepsin was the most potent compound in our screen *in vitro* (IC_50_ ≈ 3.5 nM), but it did not improve the survival rate in our metastatic mouse model (fig. 2A, 2B; fig. 3B). Romidepsin is FDA-approved for cutaneous T-cell lymphoma treatment (52) and has been shown to be potent against various other cancer types *in vitro* (53–55). *In vivo* experiments with romidepsin have been challenging, which may be attributed to its short half-life and potential long-term toxicities (56, 57). However, its high potency in UM cells highlights class I HDAC inhibition specifically as a potential vulnerability in UM, and may warrant further studies with different treatment paradigms and delivery systems to identify an applicable therapeutic window.

BET inhibition, on the other hand, has been less explored for UM treatment. While JQ-1 has demonstrated *in vitro* efficacy uveal melanoma cells, it is not tested clinically due to its short half-life, though its analogues may be more promising due to their enhanced pharmacokinetic properties (58–60). Clinical trials with BET-inhibitors PLX51107 (NCT02683395) and PLX2853 (NCT03297424), which included UM patients, both had limited success (60, 61),(62). However, here we show that different BET inhibitors have varying efficacies for UM. Our initial panel of compounds included 35 BET inhibitors, most of which did not significantly reduce the viability of UM cells (fig. 2C). Notably, JQ-1, and several of its analogues (MS417, (R)-(-)-JQ1 Enantiomer, birabresib, molibresib, (+)-JQ1 PA, JQ-1 carboxylic acid) were not efficacious, while two JQ-1 analogues, CPI-203 and BET bromodomain inhibitor (CAS: 1505453-59-7), significantly reduced UM cell viability (fig. 2C). This demonstrates a high level of diversity in compounds within the same drug class, even amongst the analogues of the same compound, and may indicate the involvement of off-target effects.

We find that the BET inhibitor mivebresib has exceptionally low toxicity towards normal fibroblasts and increases the median overall survival time by 50% *in vivo*, from 84 to 121 days, in a metastatic UM mouse model (fig. 2A, 3B). Mivebresib is an oral, small-molecule pan-BET inhibitor that induces cell death and tumor regression in animal models of malignancies such as myeloid leukemia (63), prostate cancer (64), and small cell lung cancer (65). In a clinical trial for patients with solid tumors that included 10 UM patients, mivebresib prevented tumor growth and reduced tumor volumes in a subset of UM patients (66). While these results were derived from a small cohort, they highlight, in combination with our findings, that treatment with specific BET inhibitors may be a clinically feasible option for UM. Mivebresib prevented the development of detectable spinal cord and femur metastases (fig. 3C). Bone metastasis occurs in approximately 16% of the patients with metastatic UM. While spinal cord metastases are rare (1%), brain metastases are more frequent (5%) (67, 68). Although we did not observe frequent brain metastases in our UM model, the blood-spinal cord barrier (BSCB) is similar to the blood-brain barrier (BBB) in function and morphology, potentially indicating that mivebresib may be able to cross the BSCB/BBB more efficiently than the HDAC inhibitors tested (69, 70).

Despite the high *in vitro* efficacy of HDAC inhibitors quisinostat and romidepsin, these compounds were less effective in reducing metastases and improving survival in a UM mouse model than the BET inhibitor mivebresib (fig. 3B). To illucidate the mechanisms of action of these compounds, we examined the gene expression changes they induced in UM cells. While each compound elicited unique gene expression signatures, we identified a significant overlap in the gene expression changes and enriched pathways induced by HDAC and BET inhibition. We found that HDAC inhibition led to the upregulation of PRC1 and PRC2 target genes, whereas BET inhibition acts through other targets, such as retinoic acid-related pathways (fig. 5E, 5F). While promoting cell death, we found that both HDAC and BET inhibition initially cause a phenotypic switch, reversing the clinical class 2 (high metastatic risk) gene expression signature (fig. 5A, 5B). The specific reversal of these key markers, both up- and downregulated, shows that both drug classes act by initially pushing tumor cells towards a less aggressive class 1 phenotype, rather than being generically toxic. Previous studies have demonstrated that neural progenitor cells treated with HDAC or BET inhibitors favor a neuronal over glial lineage (71–73). We similarly found that genes associated with glial and melanocytic cells were downregulated, while key neuronal genes and pathways were upregulated (fig. 6). These data indicate that, given the shared developmental neural crest origin of melanocytes and some neuronal cell types (74), the stem-like features of UM cells (75) may allow them to be pharmacologically pushed towards a neuronal phenotype.

In summary, our data reveal different mechanisms by which HDAC and BET inhibitors reduce the viability of UM cells, and highlight BET inhibition with specific BET inhibitors as a promising treatment angle for metastatic UM.

## METHODS

### Cell culture

UM (MP41, MP46, and MP38) cell line stocks were obtained from the American Type Tissue Collection (ATCC). UM cells were cultured at 37°C under normoxic conditions (5.0% CO_2_, 5% O_2_) in D-MEM/F-12 medium with 10% heat-inactivated FBS, 2 mmol/L GlutaMAX, 1 mmol/L Non-Essential Amino Acid (NEAA) cell culture supplement, 0.5× Insulin-Transferin-Selenium (ITS), and 1x Pen-Strep (10 000 U/mL, Gibco). All UM cell lines were verified using short tandem repeat (STR) analysis.

### Compound screening

For the primary screening, we tested a 932-compound epigenetic library (TargetMol, L1200) consisting of inhibitors and activators of epigenetic-modifying enzymes (writers, erasers, and readers). All stock compounds were dissolved in 100% DMSO and tested in duplicates at a test concentration of 1 μM drug and final DMSO concentration of 0.1%. Wells with Hank’s Balanced Salt Solution (HBSS) assay buffer containing 0.1% DMSO served as negative controls. Velcade (1 μM bortezomib) served as the positive control. One thousand cells per well were seeded in 384-well white microtiter plates in a humidified incubator at 37°C with 5% O_2_ and 5% CO_2_ overnight (∼16 h). The cells were then treated with compounds for 72 h. Cell viability was assessed by measuring ATP levels using a luminescence-based assay (CellTiter-Glo, Promega) on a Perkin Elmer Envision Multilabel Plate Reader. Positive hits were defined as compounds that showed cell death higher than the mean of the negative controls plus 3 standard deviations. Assays on each plate were considered valid only when the Z’-factor of the plate was equal to or greater than 0.5 (Z’ ≥ 0.5).

### Concentration-response testing

Cell lines were treated using a 10-point 1:3 dilution series starting at a nominal test concentration of 10 μM for all drugs, except romidepsin, for which the starting concentration was 300 nM (n = 4, 20 000-fold concentration range). Cell viability was assessed after 72 h of treatment by measuring ATP levels using a luminescence-based assay (CellTiter-Glo, Promega) on a Perkin Elmer Envision Multilabel Plate Reader, and normalized to the viability of cells treated with 0.1% DMSO, which served as the negative control. Four-parameter curve fitting (non-linear regression, log(inhibitor) vs. response, variable slope) performed using GraphPad Prism to measure the efficacy (% cell viability) and potency (IC_50_) of each compound.

### Animal studies

The University of Miami Institutional Animal Care and Use Committee (IACUC) approved all animal procedures. Female NOD Scid Gamma (NSG) mice were obtained from Jackson Laboratory (Stock No. 002374) and bred in-house for one generation. MP41 cells were transduced with retroviruses expressing RFP-luciferase (pMSCV-IRES-luciferase-RFP), and successful transduction was confirmed by imaging the cells on a cell imager (Zoe, Bio-Rad, Hercules, CA, USA) with an RFP filter. After transduction, RFP-positive cells were sorted and purified using fluorescence activated cell sorting (FACS). For the model generation, 1 × 10^5^ cells were injected intravenously (tail vein) into 16-week-old female NSG mice (n = 10 per group). Seven days after cell injection, drug treatments were initiated. Treatment groups assignments were randomized. Optimal drug doses were determined through prior toxicity assays, which were 2 mg/kg of romidepsin (TargetMol, T6006) via weekly intraperitoneal (IP) injection, 5 mg/kg of quisinostat (TargetMol, T6055) five times per week via IP injection, and 2 mg/kg of mivebresib (TargetMol, T3712) five times per week via oral gavage. The development of tumor metastases was monitored weekly during the course of the experiment using an in vivo imaging system (IVIS Spectrum, Revvity). Briefly, 10 min prior to imaging, mice were injected intraperitoneally with d-luciferin (Perkin Elmer, 760504) at a dose of 150 mg/kg. Mice were sacrificed at the endpoint (defined as more than 20% weight loss or significant changes in health status), and tumor metastases in different organs were quantified *ex vivo* using IVIS. Significance testing for survival curves were conducted with the log-rank (Mantel-Cox) test.

### Isolation and resistance testing of metastatic cells in mouse livers

Tumor-bearing liver tissue was minced and incubated in collagenase Type IV solution (1x D-MEM with 400 U/mL Type IV collagenase powder (Gibco) and 0.5 μg/mL Amphotericin B solution (Sigma)) overnight at 4°C. Tumor cells from the liver were seeded in UM cell media (see above) and confirmed to be MP41 cells by RFP fluorescence. Drug resistance testing was performed using with concentration-response testing (see above), with cells extracted from liver metastases from the vehicle group serving as the control.

### RNA sequencing

For the 24-hour treatment RNA-seq analysis, 100 000 cells were seeded per well in 6-well plates in triplicates for each treatment group. After cell attachment, cells were treated with romidepsin (40 nM), quisinostat (40 nM), or mivebresib (1200 nM) at a final concentration of 0.1% DMSO. Concentrations were selected through initial testing for doses that elicited morphological changes without excessive cell death in 24 hours. Wells treated with 0.1% DMSO served as the control group. Total RNA was extracted 24 hours after treatment using the Zymo Research Quick-RNA MiniPrep kit and the samples were sequenced by BGI (Cambridge, MA, USA). All samples were sequenced with over 18 million paired-end reads (150 base pairs). The treatment group files were concatenated and analyzed using BioJupies, which utilizes *limma* powered differential expression analysis (76). Pathway analysis was performed with Metascape (77) using significantly differentially expressed genes (Adj. P < 0.05, | log_2_ FC | > 1.5) and transcription factor analysis was performed using ChIP Enrichment Analysis (ChEA) (47). Data will be available on the Gene Expression Omnibus (GEO) data repository upon publication.

### iLINCS analysis

To compare the transcriptomic changes caused by our drugs to other perturbations, we used the Library of Integrated Network-based Cellular Signatures (iLINCS) (40) data portal to identify genes dysregulated by HDAC treatments. We identified 180 genes that were consistently up- or down-regulated (Adj. P < 0.05, | log_2_ FC | > 1.5) as a result of treatment with 8 different HDAC treatments (trichostatin A, vorinostat, panobinostat, dacinostat, romidepsin, belinostat, entinostat, mocetinostat) across analyzed cell lines, and determined the gene expression shifts of these genes as a result of HDAC and BET inhibitor treatment in our cell lines (fig. 4E, supplemental fig. 4E). We additionally used the connected perturbations analysis function of iLINCS to identify compounds eliciting gene signatures similar to those in our study using lists of significantly differentially expressed genes for each treatment group (Adj. P < 0.05, | log_2_ FC | > 1.5) (fig.4F, supplemental fig. 4F). We also used the correlation matrix of gene signatures elicited by compounds in the iLINCS data base and calculated the average correlation coefficient of 7 BET inhibitor treatments (JQ-1, (-)-JQ1, (+)-JQ1, JQ1+SR1277, I-BET, I-BET151, PFI-1) with 7 HDAC inhibitor treatments (entinostat, mocetinostat, rocilinostat, pracinostat, belinostat, vorinostat, MC1568). We compared the average correlation coeffecient of HDAC and BET inhibitors with each other to their correlation with the other drug classes in the dataset (fig. 4G, 4H).

### IF of neuronal markers

20 000 MP41 cells per well were seeded in chamber slides (Lab-Tek II, 155382). After cells attached, they were treated with romidepsin (40 nM), quisinostat (40 nM), mivebresib (1200 nM), or 0.01% DMSO (control). Following 24 hours of treatment, cells were fixed with a 10 minute 4% paraformaldehyde incubation. Immunocytochemistry was performed as described in Abcam Immunocytochemistry protocol (78). The cells in each treatment group were incubated with an antibody for either Synapsin-1 (SYN1) (Cell Signaling, D12G5) or Beta-Tubulin (TUBB3) (Cell Signaling, D71G9). Alexa-Fluor anti-rabbit antibody was used for visualization (Invitrogen, A-11012). DAPI (Sigma-Aldrich, MBD0015) diluted 1:1 000 in phosphate buffered saline (PBS) was added prior to visualization. The cells were visualized in 40x on a fluorescent microscope using the Infinity Analyze program.

### Western of neuronal markers

100 000 MP41 cells were seeded per well in 6-well plates. After cells attached, they were treated with romidepsin (40 nM), quisinostat (40 nM), mivebresib (1 200 nM), or 0.01% DMSO (control). Following 24 hours of treatment, cells were pelleted and lysed with 50 μL of RIPA buffer containing protease inhibitor (Roche cOmplete ULTRA Tablet, 5892970001). Samples were sonicated and centrifuged at maximum speed (16 000g) for 15 minutes in 4°C to pellet cellular debris. The supernatants were transferred to new tubes, and protein was quantified with BCA assay (Thermo Scientific Pierce BCA Protein Assay, A55864). 50 μg of protein was boiled with Laemli buffer + BME at 1/3 of the protein sample for 5 minutes at 95°C. Protein samples were separated on precast polyacrylamide gel (4–15%) (Bio-Rad, 5678084) and transferred to nitrocellulose membrane via Trans-Blot Turbo System (Bio-Rad, 170–4159). Membrane was blocked with 5% BSA in 0.1% Tween 20 in TBS (TBS-T) for 1 h at room temperature (RT), followed by incubation with primary antibodies for Synapsin-1 (SYN1) (Cell Signaling, D12G5), beta-tubulin III (TUBB3) (Cell Signaling, D71G9), and beta-actin (ACTB) (Santa Cruz, sc-47778) diluted in 5% BSA in TBS-T overnight at 4 °C. Membranes were washed with TBS-T three times and once with TBS, then incubated in secondary antibodies (LI-COR IRDye, 926-32210, 926-68073) diluted in 5% BSA in TBS-T for 1 h at RT. The membranes were washed with TBS-T three times and once with TBS, then visualized on Odyssey CLx LI-COR imager.

## Supporting information

Supplemental Material- Uncropped Westerns

## Acknowledgements

This work was supported by funds from the Sylvester Comprehensive Cancer Center (SCCC) and Interdisciplinary Stem Cell Institute (ISCI), the American Cancer Society (ACS) Discovery Boost Grant, the Elsa Pardee Foundation, the Sinskey Foundation, and NIH NEI R21EY036185-01 (S.K.). We thank the Cancer Modeling Shared Resource (CMSR, RRID: SCR_022891) from the Sylvester Comprehensive Cancer Center (SCCC) for support with in vivo modeling, efficacy studies, noninvasive imaging, and histological work. We thank the Molecular Therapeutic Shared Resource (MTSR) of the SCCC for drug screening support. This work was also supported by funds from the National Cancer Institute (NCI) Cancer Center Support Grants P30 CA142543 to University of Texas Southwestern Simmons Comprehensive Cancer Center (J.W.H.), Cancer Prevention and Research Institute of Texas Recruitment of Established Investigator Award RR220010 (J.W.H.), the NEI Center Core Grants P30 EY030413 to University of Texas Southwestern Department of Ophthalmology (J.W.H.), and the Research to Prevent Blindness, Inc. Challenge Grant to University of Texas Southwestern Department of Ophthalmology (J.W.H.).

The Bascom Palmer Eye Institute received funding from the National Eye Institute, Grant P30 EY014801, and Research to Prevent Blindness Unrestricted Grant GR004596-1. The Sylvester Comprehensive Cancer Center received funding from the National Cancer Institute (Grant P30 CA240139).

The authors declare no conflicts of interest.

## Supplemental Figures

**Supplemental Table 1.** Primary screen results. IC_50_ (M) values of the hit compounds identified by the primary screen for each UM cell line, along with the mechanisms of action for each compound.

| Name | MP41 IC50 (M) | MP38 IC50 (M) | MP46 IC50 (M) | Average IC50 (M) | Mechanism |
| --- | --- | --- | --- | --- | --- |
| Alobresib | 1.04E-07 | 1.404E-07 | 7.122E-08 | 1.0514E-07 | BET inhibitor |
| Podofilox | 6.49E-09 | 1.503E-08 | 6.562E-09 | 9.362E-09 | Topoisomerase II inhibitor |
| Staurosporine | 2.71E-07 | 5.194E-08 | 6.863E-07 | 3.3648E-07 | PKCα, PKCγ, PKCη inhibitor |
| SKLB-23bb | 8.44E-08 | 1.742E-05 | 9.481E-08 | 5.86639E-06 | HDAC6 inhibitor |
| GSK1324726A | 1.54E-07 | 2.475E-07 | 1.045E-07 | 1.68533E-07 | BRD2, BRD3, BRD4 inhibitor |
| (S)-(+)-Camptothecin | 1.17E-07 | 6.757E-07 | 2.102E-07 | 3.342E-07 | Topoisomerase I inhibitor |
| Gemcitabine | 7.69E-08 | 1.233E-06 | 1.686E-07 | 4.92843E-07 | DNA synthesis inhibitor |
| CPI203 | 1.53E-07 | 1.604E-07 | 1.091E-07 | 1.40733E-07 | BRD4 inhibitor |
| NSC228155 | 2.59E-06 | 3.216E-07 | 2.789E-06 | 1.90087E-06 | EGFR activator |
| BET BRD Inhibitor | 3.76E-07 | 3.445E-07 | 2.477E-07 | 3.22667E-07 | BET inhibitor |
| Panobinostat | 1.43E-08 | 3.575E-08 | 2.768E-08 | 2.58967E-08 | HDAC inhibitor |
| Quisinostat | 9.12E-09 | 1.162E-08 | 1.983E-08 | 1.35243E-08 | HDAC inhibitor |
| Fimepinostat | 5.96E-09 | 1.758E-08 | 9.285E-09 | 1.09427E-08 | HDAC and PI3K inhibitor |
| Cucurbitacin B | 4.84E-08 | 1.41E-08 | 5.111E-08 | 3.78567E-08 | PI3K/AKT inhibitor |
| Romidepsin | 3.16E-14 | 8.916E-10 | 1.269E-09 | 7.20211E-10 | Class I HDAC inhibitor |
| AZD5153 | 9.04E-08 | 8.867E-08 | 6.833E-08 | 8.24767E-08 | BRD4 inhibitor |
| Mivebresib | 9.96E-08 | 1.877E-07 | 8.921E-08 | 1.2549E-07 | BET inhibitor |
| ABBV-744 | 8.127 | 2834 | 0.05655 | 947.3945167 | BRD4 inhibitor |
| Quisinostat 2HCl | 5.42E-09 | 6.754E-09 | 1.385E-08 | 8.674E-09 | HDAC inhibitor |
| 666-15 | 5.89E-05 | 0.0005853 | 0.002194 | 0.00094607 | EGFR inhibitor |
| Velcade | 4.34E-09 | 1.487E-08 | 3.519E-09 | 7.57633E-09 | Proteasome inhibitor |

**Supplemental Figure 1.**
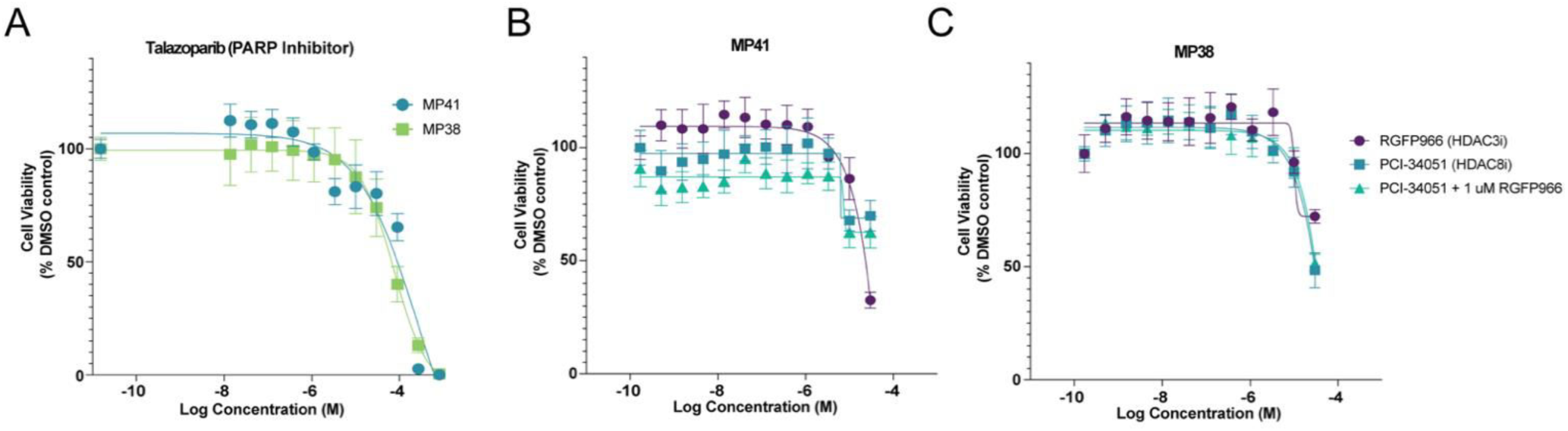
PARP inhibitor, HDAC3 inhibitor, and HDAC8 inhibitor concentration-response testing. **(A)** Concentration-response curves of MP41 and MP38 cells treated with the PARP inhibitor talazoparib. Error bars indicate SD. n=4 for each concentration. **(B)** Concentration-response curves of MP41 and MP38 cells treated with HDAC3 (RGFP966) and HDAC8 (PCI-34051) inhibitors. Error bars indicate SD. n=4 for each concentration.

**Supplemental Figure 2.**
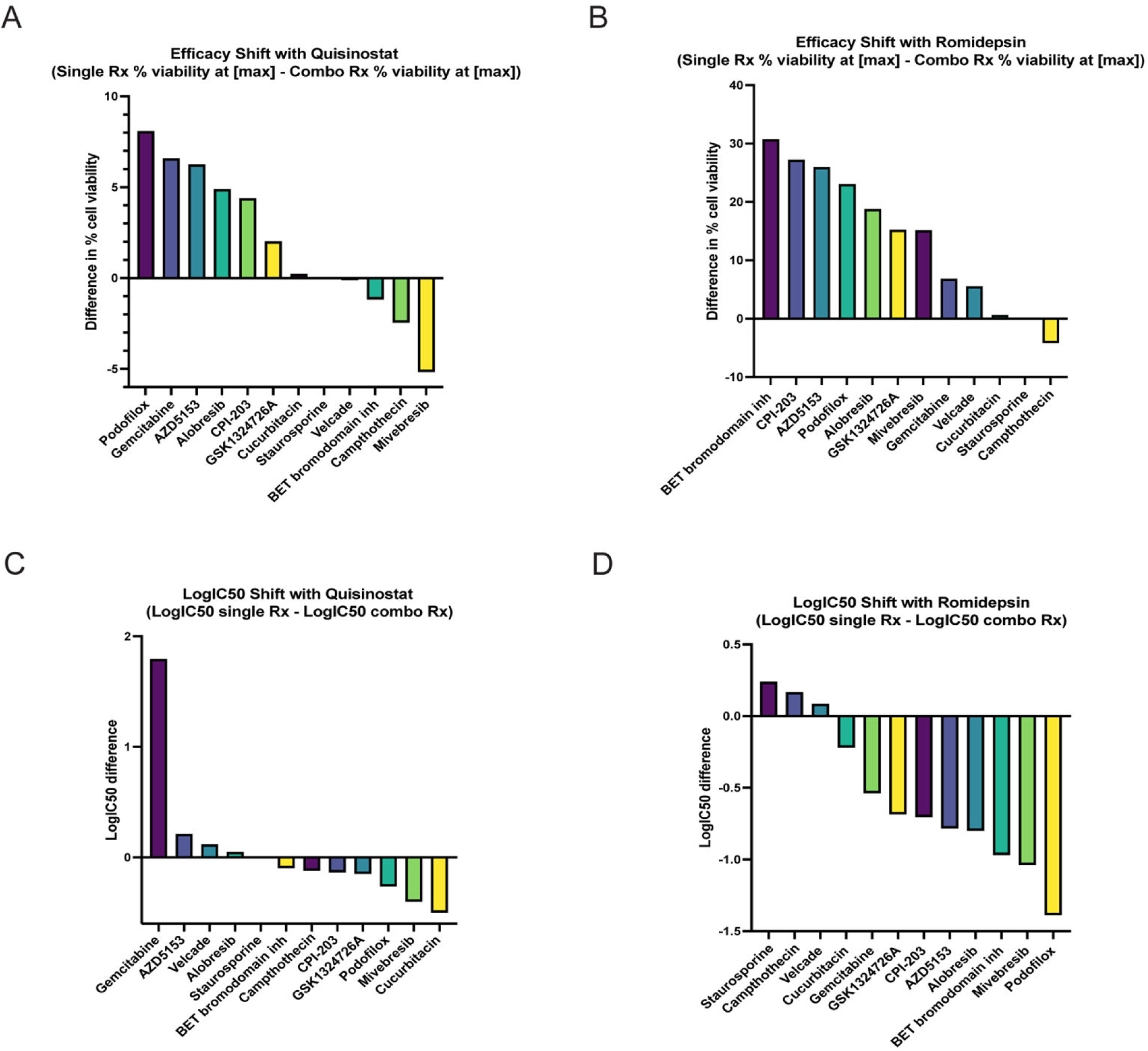
Synergistic tests of quisinostat and romidepsin with other candidate compounds. **(A)** Difference in percent cell viability at the highest concentration (10 µM) for cells treated with quisinostat plus EC_20_ of other candidate compound relative to cell viability when treated with only 10 µM quisinostat. Greater positive values indicate better synergy. **(B)** Difference in percent cell viability at the highest concentration (10 µM) for cells treated with romidepsin plus EC_20_ of other candidate compounds relative to cell viability when treated with only 10 µM romidepsin. Greater positive values indicate better synergy. **(C)** Log IC_50_ shift of cells treated with Quisinostat and the EC_20_ of other candidate compounds relative to cells treated with only quisinostat. Greater positive values indicate better synergy. **(D)** Log IC_50_ shift of cells treated with romidepsin and the EC_20_ of other candidate compound relative to cells treated with only romidepsin. Greater positive values indicate better synergy.

**Supplemental Figure 3.**
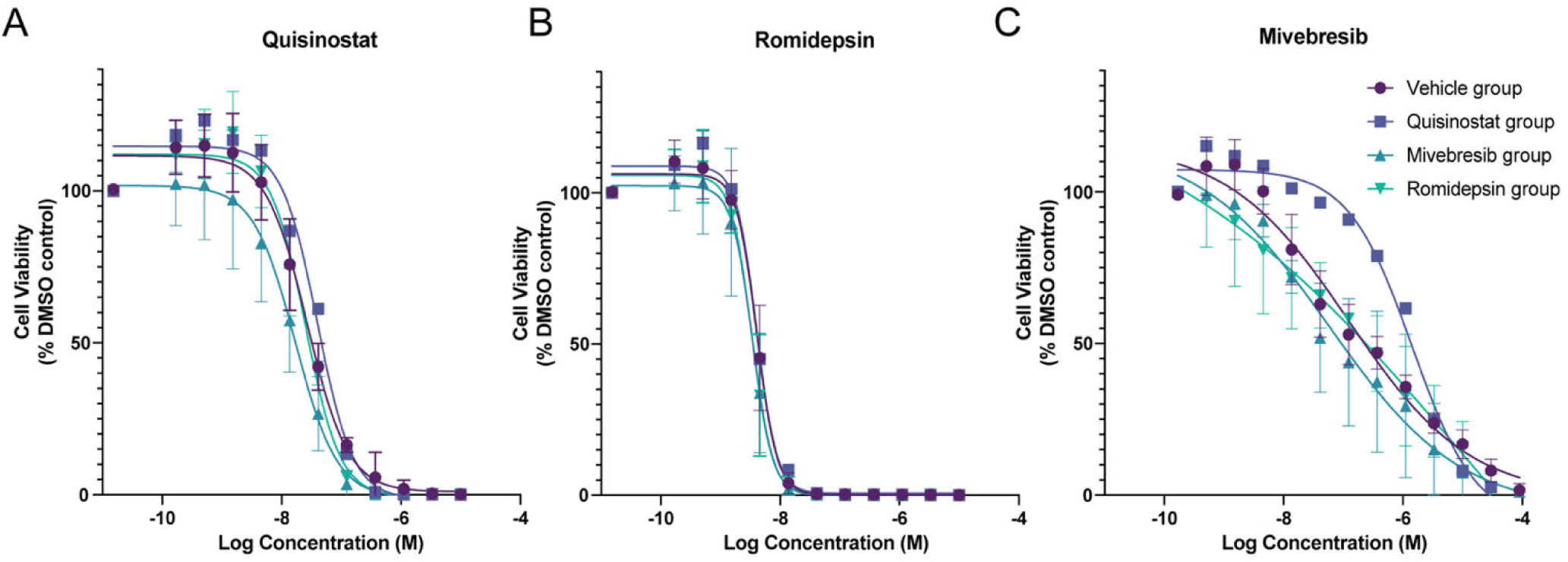
E*x vivo* testing of acquired drug resistance in vehicle-and treated tumor cells from murine livers. **(A)** Concentration-response curves for quisinostat treatment of MP41 cells extracted from mouse liver tumor samples (vehicle n =3; quisinostat n =1, mivebresib n = 4, romidepsin n = 3). Error bars indicate SD. **(B**) Concentration-response curves for romidepsin treatment of MP41 cells extracted from mouse liver tumor samples (vehicle n =3; quisinostat n =1, mivebresib n = 4, romidepsin n = 3). Error bars indicate SD. **(C)** Concentration-response curve of mivebresib treatment of MP41 cells extracted from mouse liver tumor samples (vehicle n =3; quisinostat n =1, mivebresib n = 4, romidepsin n = 3). Error bars indicate SD.

**Supplemental Figure 4.**
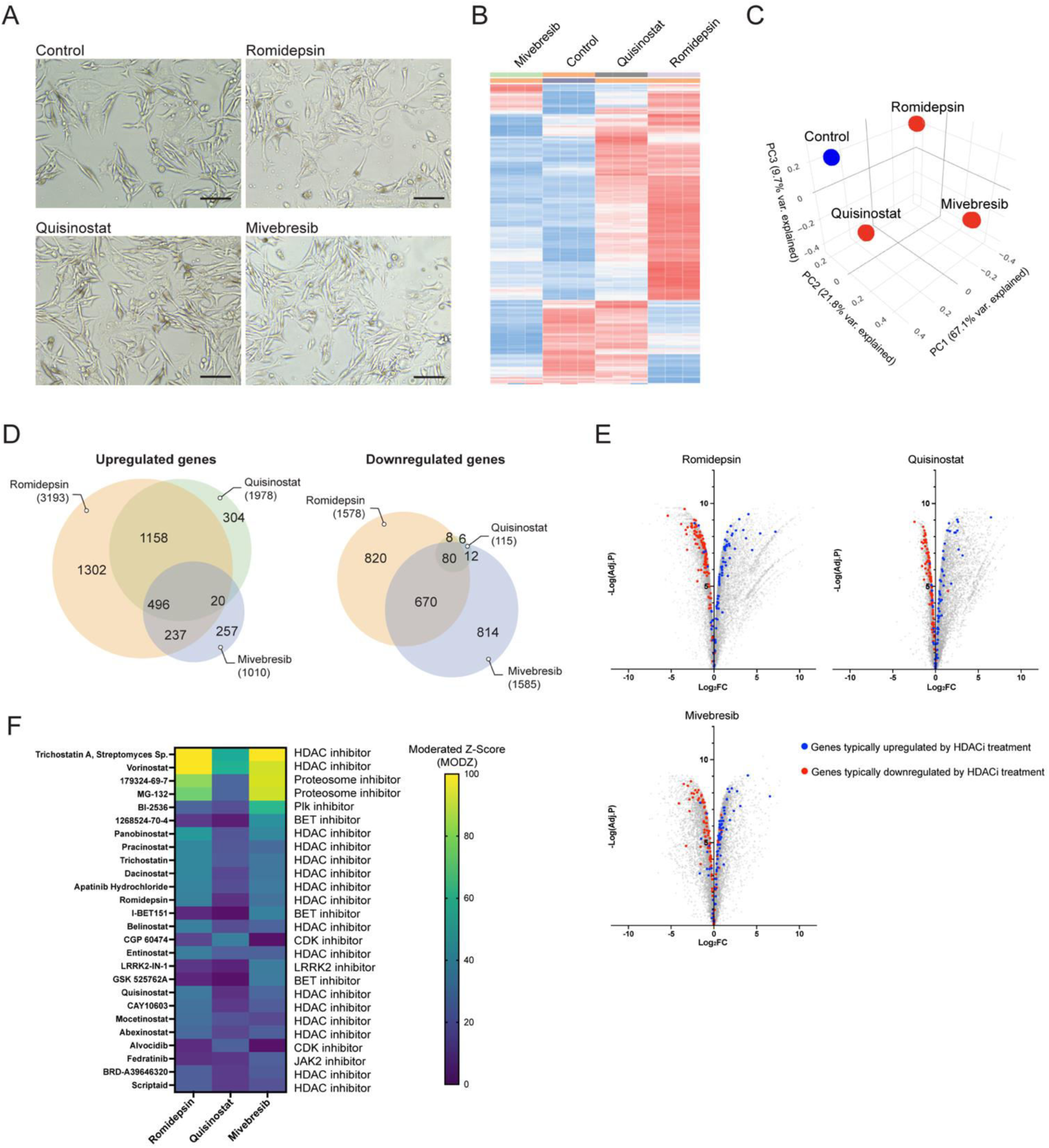
RNA-seq analysis of MP46 cells treated with candidate compounds for 24 h. **(A)** Images of MP46 cells treated with each compound for 24 hours. Scale bar = 100 µm. **(B)** Heatmap clustering of changes in gene expression of MP46 cells per treatment group (n = 3 per condition). **(C)** PCA clustering of replicates for each treatment in MP46 cells. **(D)** Venn diagram depicting overlaps between the treatment groups of significantly upregulated and downregulated genes in drug-treated MP46 cells (Adj. P < 0.05, | log_2_ FC | > 1.5).. **(E)** Volcano plot of changes in gene expression relative to the control for each treatment group in MP46 cells. Blue and red dots are 180 genes found to be consistently dysregulated as a result of eight HDAC inhibitor treatments in iLINCS. Blue dots are genes that were consistently upregulated by HDAC inhibitor treatment (n = 77), while red dots are genes that were consistently downregulated (n=103). **(F)** Heatmap of moderated Z-score (MODZ) of compounds inducing similar gene expression signatures to romidepsin, quisinostat, and mivebresib treatment in MP46 cells using iLINCS connected perturbation analysis. Higher MODZ indicates greater similarity.

**Supplemental Figure 5.**
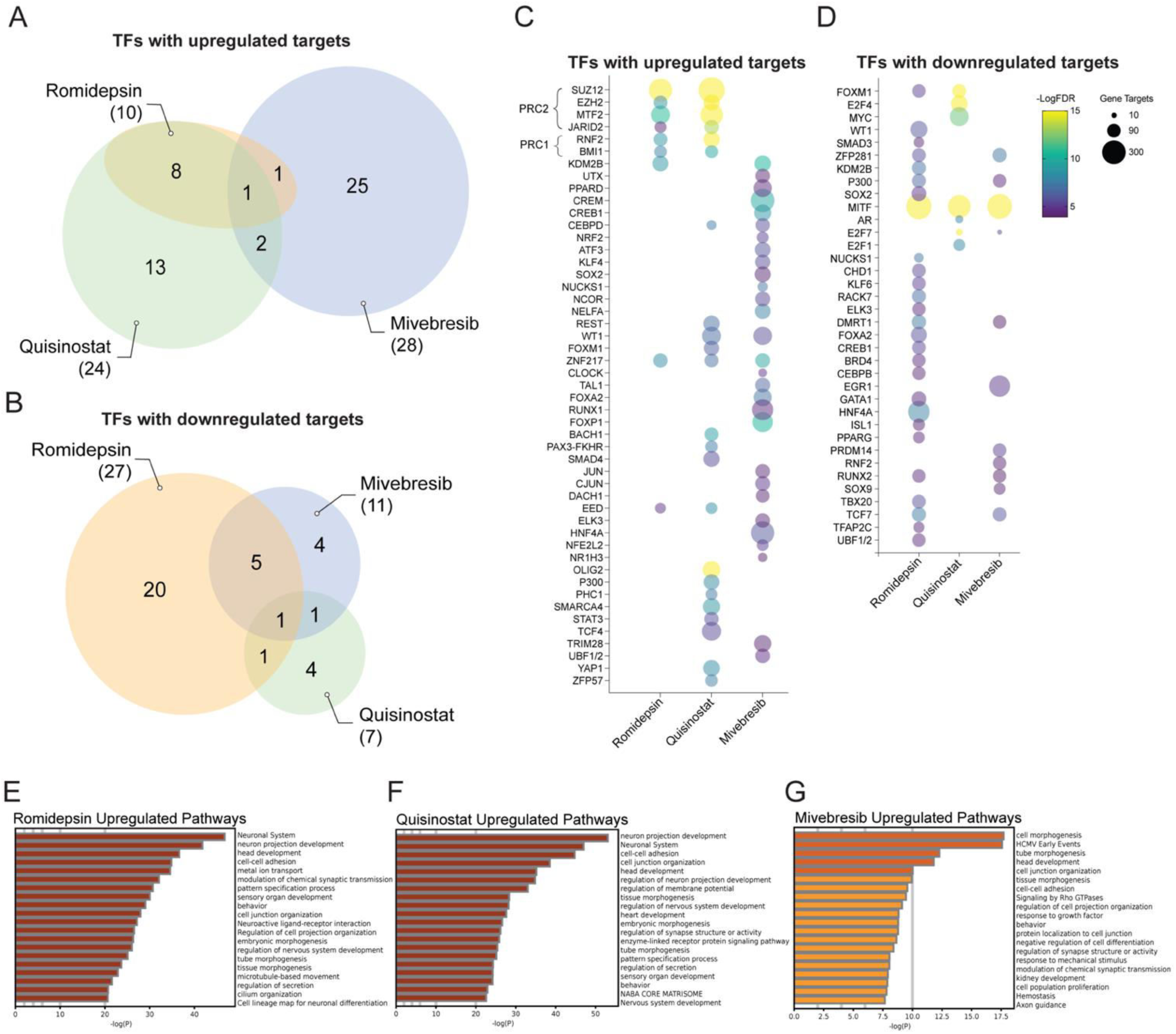
BET and HDAC inhibition mechanisms and pathway changes in MP46 cells. **(A)** Venn diagram showing overlaps in predicted transcription factors with upregulated and **(B)** downregulated gene targets, inferred by gene expression changes induced by each treatment in MP46 cells.. **(C)** Bubble plot of the top predicted transcription factors with upregulated and **(D)** downregulated gene targets for each treatment in MP46 cells. Color scheme indicates -Log_10_FDR of each predicted transcription factor, and bubble size is determined by the number of corresponding gene targets. **(E-G)** Upregulated pathways in drug-treated MP46 cells predicted from list of significantly upregulated genes in each treatment group (log_2_ FC > 1.5, adj. p < 0.05).

**Supplemental Figure 6.**
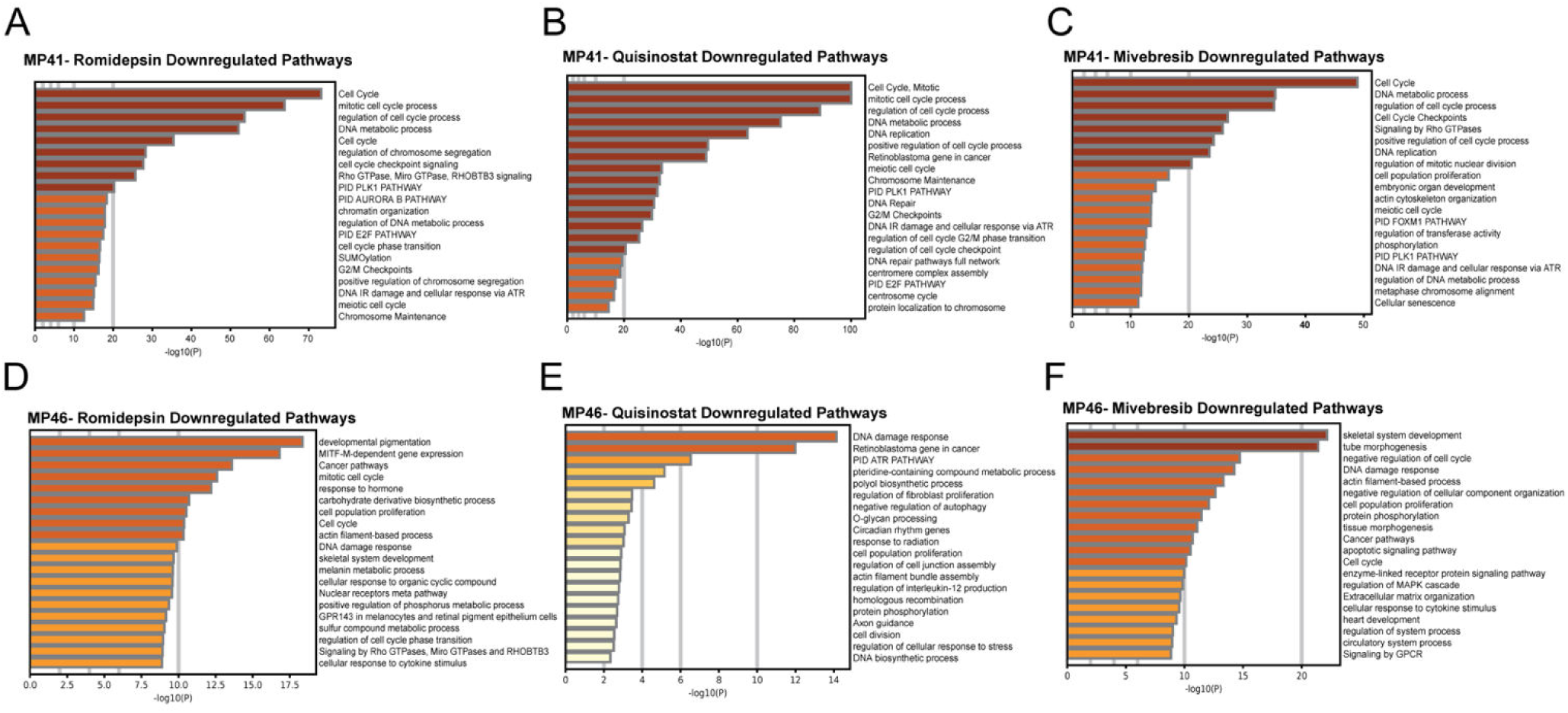
Pathways downregulated by each drug in MP41 and MP46 cells. **(A-C)** Downregulated pathways in drug-treated MP41 cells predicted from list of significantly downregulated genes in each treatment group (log_2_ FC < −1.5, adj. P value < 0.05). **(D-F)** Downregulated pathways in drug-treated MP46 cells predicted from list of significantly downregulated genes in each treatment group (log_2_ FC < −1.5, adj. P value < 0.05).

## References

1. Carvajal RD, Sacco JJ, Jager MJ, Eschelman DJ, Olofsson Bagge R, Harbour JW, et al. Advances in the clinical management of uveal melanoma. Nature Reviews Clinical Oncology. 2023;20(2):99–115.

2. Nathan P, Hassel JC, Rutkowski P, Baurain J-F, Butler MO, Schlaak M, et al. Overall survival benefit with tebentafusp in metastatic uveal melanoma. New England Journal of Medicine. 2021;385(13):1196–206.

3. Decatur CL, Ong E, Garg N, Anbunathan H, Bowcock AM, Field MG, et al. Driver mutations in uveal melanoma: associations with gene expression profile and patient outcomes. JAMA ophthalmology. 2016;134(7):728–33.

4. Van Raamsdonk CD, Bezrookove V, Green G, Bauer J, Gaugler L, O’Brien JM, et al. Frequent somatic mutations of GNAQ in uveal melanoma and blue naevi. Nature. 2009;457(7229):599–602.

5. Van Raamsdonk CD, Griewank KG, Crosby MB, Garrido MC, Vemula S, Wiesner T, et al. Mutations in GNA11 in uveal melanoma. New England Journal of Medicine. 2010;363(23):2191–9.

6. Johansson P, Aoude LG, Wadt K, Glasson WJ, Warrier SK, Hewitt AW, et al. Deep sequencing of uveal melanoma identifies a recurrent mutation in PLCB4. Oncotarget. 2016;7(4):4624.

7. Moore AR, Ceraudo E, Sher JJ, Guan Y, Shoushtari AN, Chang MT, et al. Recurrent activating mutations of G-protein-coupled receptor CYSLTR2 in uveal melanoma. Nature genetics. 2016;48(6):675–80.

8. Onken MD, Worley LA, Long MD, Duan S, Council ML, Bowcock AM, et al. Oncogenic Mutations in GNAQ Occur Early in Uveal Melanoma. Investigative Opthalmology & Visual Science. 2008;49(12):5230.

9. Vader M, Madigan M, Versluis M, Suleiman H, Gezgin G, Gruis NA, et al. GNAQ and GNA11 mutations and downstream YAP activation in choroidal nevi. British journal of cancer. 2017;117(6):884–7.

10. Harbour JWO, Michael D; Roberson, Elisha D O; Duan, Shenghui; Cao, Li; Worley, Lori A; Council, M Laurin; Matatal, Katie A; Helms, Cynthia; Bowcock, Anne M Frequent Mutation of BAP1 in Metastasizing Uveal Melanomas. SCIENCE. 2010;330(6009):1410–3.

11. Harbour JW, Roberson EDO, Anbunathan H, Onken MD, Worley LA, Bowcock AM. Recurrent mutations at codon 625 of the splicing factor SF3B1 in uveal melanoma. Nature Genetics. 2013;45(2):133–5.

12. Martin M, Maßhöfer L, Temming P, Rahmann S, Metz C, Bornfeld N, et al. Exome sequencing identifies recurrent somatic mutations in EIF1AX and SF3B1 in uveal melanoma with disomy 3. Nature genetics. 2013;45(8):933–6.

13. Durante MA, Field MG, Sanchez MI, Covington KR, Decatur CL, Dubovy SR, et al. Genomic evolution of uveal melanoma arising in ocular melanocytosis. Molecular Case Studies. 2019;5(4):a004051.

14. Durante MA, Rodriguez DA, Kurtenbach S, Kuznetsov JN, Sanchez MI, Decatur CL, et al. Single-cell analysis reveals new evolutionary complexity in uveal melanoma. Nature Communications. 2020;11(1).

15. Field MG, Durante MA, Anbunathan H, Cai LZ, Decatur CL, Bowcock AM, et al. Punctuated evolution of canonical genomic aberrations in uveal melanoma. Nature Communications. 2018;9(1).

16. Campagne A, Lee M-K, Zielinski D, Michaud A, Le Corre S, Dingli F, et al. BAP1 complex promotes transcription by opposing PRC1-mediated H2A ubiquitylation. Nature communications. 2019;10(1):348.

17. Yu H, Mashtalir N, Daou S, Hammond-Martel I, Ross J, Sui G, et al. The ubiquitin carboxyl hydrolase BAP1 forms a ternary complex with YY1 and HCF-1 and is a critical regulator of gene expression. Molecular and cellular biology. 2010;30(21):5071–85.

18. Field MK, Jeffim N; Bussies, Parker L; Cai, Louie Z; Alawa, Karam A; Decatur, Christina L; Kurtenbach, Stefan; Harbour, J William. BAP1 Loss Is Associated with DNA Methylomic Repatterning in Highly Aggressive Class 2 Uveal Melanomas. Clinical cancer research. 2019;25(18):5663.

19. Kuznetsov JN, Aguero TH, Owens DA, Kurtenbach S, Field MG, Durante MA, et al. BAP1 regulates epigenetic switch from pluripotency to differentiation in developmental lineages giving rise to BAP1-mutant cancers. Science advances. 2019;5(9):eaax1738.

20. Bakhoum MF, Francis JH, Agustinus A, Earlie EM, Di Bona M, Abramson DH, et al. Loss of polycomb repressive complex 1 activity and chromosomal instability drive uveal melanoma progression. Nature communications. 2021;12(1):5402.

21. Carbone M, Harbour JW, Brugarolas J, Bononi A, Pagano I, Dey A, et al. Biological mechanisms and clinical significance of BAP1 mutations in human cancer. Cancer discovery. 2020;10(8):1103–20.

22. Némati F, Sastre-Garau X, Laurent C, Couturier J, Mariani P, Desjardins L, et al. Establishment and characterization of a panel of human uveal melanoma xenografts derived from primary and/or metastatic tumors. Clinical cancer research. 2010;16(8):2352–62.

23. Adams J, Kauffman M. Development of the proteasome inhibitor Velcade™(Bortezomib). Cancer investigation. 2004;22(2):304–11.

24. Schmittel A, Schmidt-Hieber M, Martus P, Bechrakis N, Schuster R, Siehl J, et al. A randomized phase II trial of gemcitabine plus treosulfan versus treosulfan alone in patients with metastatic uveal melanoma. Annals of oncology. 2006;17(12):1826–9.

25. Lapadula D, Farias E, Randolph CE, Purwin TJ, McGrath D, Charpentier TH, et al. Effects of oncogenic Gαq and Gα11 inhibition by FR900359 in uveal melanoma. Molecular Cancer Research. 2019;17(4):963–73.

26. Liu LF, Desai SD, LI TK, Mao Y, Sun M, SIM SP. Mechanism of action of camptothecin. Annals of the New York Academy of Sciences. 2000;922(1):1–10.

27. Gardner TJ, Cohen T, Redmann V, Lau Z, Felsenfeld D, Tortorella D. Development of a high-content screen for the identification of inhibitors directed against the early steps of the cytomegalovirus infectious cycle. Antiviral research. 2015;113:49–61.

28. Garg S, Kaul SC, Wadhwa R. Cucurbitacin B and cancer intervention: Chemistry, biology and mechanisms. International journal of oncology. 2018;52(1):19–37.

29. Landreville SA, Olga A; Matatall, Katie A; Kneass, Zachary T; Onken, Michael D; Lee, Ryan S; Bowcock, Anne M; Harbour, J William. Histone Deacetylase Inhibitors Induce Growth Arrest and Differentiation in Uveal Melanoma. Clinical cancer research. 2012;18(2):408.

30. Kuznetsoff JNO, Dawn A.; Lopez, Andy; Rodriguez, Daniel A.; Chee, Nancy T.; Kurtenbach, Stefan; Bilbao, Daniel; Roberts, Evan R.; Volmar, Claude-Henry; Wahlestedt, Claes; Brothers, Shaun P.; Harbour, J. William. Dual Screen for Efficacy and Toxicity Identifies HDAC Inhibitor with Distinctive Activity Spectrum for BAP1-Mutant Uveal Melanoma. Molecular cancer research. 2021;19(2):215.

31. Moschos MM, Dettoraki M, Androudi S, Kalogeropoulos D, Lavaris A, Garmpis N, et al. The role of histone deacetylase inhibitors in uveal melanoma: current evidence. Anticancer research. 2018;38(7):3817–24.

32. Wang Y, Liu M, Jin Y, Jiang S, Pan J. In vitro and in vivo anti-uveal melanoma activity of JSL-1, a novel HDAC inhibitor. Cancer letters. 2017;400:47–60.

33. Dai W, Zhou J, Jin B, Pan J. Class III-specific HDAC inhibitor Tenovin-6 induces apoptosis, suppresses migration and eliminates cancer stem cells in uveal melanoma. Scientific reports. 2016;6(1):22622.

34. Nicolas E, Yamada T, Cam HP, FitzGerald PC, Kobayashi R, Grewal SI. Distinct roles of HDAC complexes in promoter silencing, antisense suppression and DNA damage protection. Nature structural & molecular biology. 2007;14(5):372–80.

35. Witt O, Deubzer HE, Milde T, Oehme I. HDAC family: What are the cancer relevant targets? Cancer letters. 2009;277(1):8–21.

36. Gentien D, Saberi-Ansari E, Servant N, Jolly A, de la Grange P, Némati F, et al. Multi-omics comparison of malignant and normal uveal melanocytes reveals molecular features of uveal melanoma. Cell reports. 2023;42(9).

37. Slaughter MJ, Shanle EK, Khan A, Chua KF, Hong T, Boxer LD, et al. HDAC inhibition results in widespread alteration of the histone acetylation landscape and BRD4 targeting to gene bodies. Cell reports. 2021;34(3).

38. Dhalluin C, Carlson JE, Zeng L, He C, Aggarwal AK, Zhou M-M, et al. Structure and ligand of a histone acetyltransferase bromodomain. Nature. 1999;399(6735):491–6.

39. Yang Z, Yik JH, Chen R, He N, Jang MK, Ozato K, et al. Recruitment of P-TEFb for stimulation of transcriptional elongation by the bromodomain protein Brd4. Molecular cell. 2005;19(4):535–45.

40. Pilarczyk M, Fazel-Najafabadi M, Kouril M, Shamsaei B, Vasiliauskas J, Niu W, et al. Connecting omics signatures and revealing biological mechanisms with iLINCS. Nature communications. 2022;13(1):4678.

41. Onken MD, Worley LA, Ehlers JP, Harbour JW. Gene expression profiling in uveal melanoma reveals two molecular classes and predicts metastatic death. Cancer research. 2004;64(20):7205–9.

42. Harbour JW. A prognostic test to predict the risk of metastasis in uveal melanoma based on a 15-gene expression profile. Molecular Diagnostics for Melanoma: Methods and Protocols. 2014:427–40.

43. Harbour JW, Chen R. The DecisionDx-UM gene expression profile test provides risk stratification and individualized patient care in uveal melanoma. PLoS currents. 2013;5.

44. Kurtenbach S, Sanchez MI, Kuznetsoff J, Rodriguez DA, Weich N, Dollar JJ, et al. PRAME induces genomic instability in uveal melanoma. Oncogene. 2023:1–11.

45. Field MG, Decatur CL, Kurtenbach S, Gezgin G, Van Der Velden PA, Jager MJ, et al. PRAME as an independent biomarker for metastasis in uveal melanoma. Clinical cancer research. 2016;22(5):1234–42.

46. Harbour JW, Correa ZM, Schefler AC, Mruthyunjaya P, Materin MA, Aaberg TA, Jr., et al. 15-Gene Expression Profile and PRAME as Integrated Prognostic Test for Uveal Melanoma: First Report of Collaborative Ocular Oncology Group Study No. 2 (COOG2.1). J Clin Oncol. 2024:JCO2400447.

47. Lachmann A, Xu H, Krishnan J, Berger SI, Mazloom AR, Ma’ayan A. ChEA: transcription factor regulation inferred from integrating genome-wide ChIP-X experiments. Bioinformatics. 2010;26(19):2438–44.

48. Bai X, Li S, Luo Y. FOXM1 promote the growth and metastasis of uveal melanoma cells by regulating CDK2 expression. International Ophthalmology. 2024;44(1):55.

49. Quintanilla RA, Utreras E, Cabezas-Opazo FA. Role of PPARγ in the Differentiation and Function of Neurons. PPAR research. 2014;2014(1):768594.

50. Simandi Z, Horvath A, Cuaranta-Monroy I, Sauer S, Deleuze J-F, Nagy L. RXR heterodimers orchestrate transcriptional control of neurogenesis and cell fate specification. Molecular and cellular endocrinology. 2018;471:51–62.

51. Schmidt A, Vogel R, Holloway MK, Rutledge SJ, Friedman O, Yang Z, et al. Transcription control and neuronal differentiation by agents that activate the LXR nuclear receptor family. Molecular and cellular endocrinology. 1999;155(1-2):51–60.

52. VanderMolen KM, McCulloch W, Pearce CJ, Oberlies NH. Romidepsin (Istodax, NSC 630176, FR901228, FK228, depsipeptide): a natural product recently approved for cutaneous T-cell lymphoma. The Journal of antibiotics. 2011;64(8):525–31.

53. Mayr C, Kiesslich T, Erber S, Bekric D, Dobias H, Beyreis M, et al. HDAC screening identifies the HDAC class I inhibitor romidepsin as a promising epigenetic drug for biliary tract cancer. Cancers. 2021;13(15):3862.

54. Panicker J, Li Z, McMahon C, Sizer C, Steadman K, Piekarz R, et al. Romidepsin (FK228/depsipeptide) controls growth and induces apoptosis in neuroblastoma tumor cells. Cell cycle. 2010;9(9):1830–8.

55. Li L-H, Zhang P-R, Cai P-Y, Li Z-C. Histone deacetylase inhibitor, Romidepsin (FK228) inhibits endometrial cancer cell growth through augmentation of p53-p21 pathway. Biomedicine & Pharmacotherapy. 2016;82:161–6.

56. Rivers ZT, Oostra DR, Westholder JS, Vercellotti GM. Romidepsin-associated cardiac toxicity and ECG changes: A case report and review of the literature. Journal of Oncology Pharmacy Practice. 2018;24(1):56–62.

57. Klimek VM, Fircanis S, Maslak P, Guernah I, Baum M, Wu N, et al. Tolerability, pharmacodynamics, and pharmacokinetics studies of depsipeptide (romidepsin) in patients with acute myelogenous leukemia or advanced myelodysplastic syndromes. Clinical Cancer Research. 2008;14(3):826–32.

58. Liu W, Cui Z, Wan Q, Liu Y, Chen M, Cheng Y, et al. The BET inhibitor JQ1 suppresses tumor survival by ABCB5-mediated autophagy in uveal melanoma. Cellular Signalling. 2025;125:111483.

59. Chen X, Huang R, Zhang Z, Song X, Shen J, Wu Q. Bet bromodomain inhibition potentiates ocular melanoma therapy by inducing cell cycle arrest. Investigative Ophthalmology & Visual Science. 2024;65(8):11-.

60. Croce M, Ferrini S, Pfeffer U, Gangemi R. Targeted therapy of uveal melanoma: Recent failures and new perspectives. Cancers. 2019;11(6):846.

61. Patnaik A, Carvajal RD, Komatsubara KM, Britten CD, Wesolowski R, Michelson G, et al. Phase ib/2a study of PLX51107, a small molecule BET inhibitor, in subjects with advanced hematological malignancies and solid tumors. American Society of Clinical Oncology; 2018.

62. Liu Xl, Run-hua Z, Pan Jx, Li Zj, Yu L, Li Yl. Emerging therapeutic strategies for metastatic uveal melanoma: Targeting driver mutations. Pigment Cell & Melanoma Research. 2024;37(3):411–25.

63. Albert DH, Goodwin NC, Davies AM, Rowe J, Feuer G, Boyiadzis M, et al. Co-clinical modeling of the activity of the BET inhibitor mivebresib (ABBV-075) in AML. in vivo. 2022;36(4):1615–27.

64. Faivre EJ, Wilcox D, Lin X, Hessler P, Torrent M, He W, et al. Exploitation of castration-resistant prostate cancer transcription factor dependencies by the novel BET inhibitor ABBV-075. Molecular Cancer Research. 2017;15(1):35–44.

65. Lam LT, Lin X, Faivre EJ, Yang Z, Huang X, Wilcox DM, et al. Vulnerability of small-cell lung cancer to apoptosis induced by the combination of BET bromodomain proteins and BCL2 inhibitors. Molecular cancer therapeutics. 2017;16(8):1511–20.

66. Piha-Paul SA, Sachdev JC, Barve M, LoRusso P, Szmulewitz R, Patel SP, et al. First-in-human study of mivebresib (ABBV-075), an oral pan-inhibitor of bromodomain and extra terminal proteins, in patients with relapsed/refractory solid tumors. Clinical Cancer Research. 2019;25(21):6309–19.

67. Group TCOMS. Assessment of Metastatic Disease Status at Death in 435 Patients With Large Choroidal Melanoma in the Collaborative Ocular Melanoma Study (COMS): COMS Report No. 15. Archives of Ophthalmology. 2001;119(5):670–6.

68. Wei AZ, Uriel M, Porcu A, Manos MP, Mercurio AC, Caplan MM, et al. Characterizing metastatic uveal melanoma patients who develop symptomatic brain metastases. Frontiers in Oncology. 2022;12:961517.

69. Sullivan JM, Badimon A, Schaefer U, Ayata P, Gray J, Chung C-w, et al. Autism-like syndrome is induced by pharmacological suppression of BET proteins in young mice. Journal of Experimental Medicine. 2015;212(11):1771–81.

70. Govindarajan V, Shah AH, Di L, Rivas S, Suter RK, Eichberg DG, et al. Systematic review of epigenetic therapies for treatment of IDH-mutant glioma. World neurosurgery. 2022;162:47–56.

71. Siebzehnrubl FA, Buslei R, Eyupoglu IY, Seufert S, Hahnen E, Blumcke I. Histone deacetylase inhibitors increase neuronal differentiation in adult forebrain precursor cells. Experimental Brain Research. 2007;176:672–8.

72. Hsieh J, Nakashima K, Kuwabara T, Mejia E, Gage FH. Histone deacetylase inhibition-mediated neuronal differentiation of multipotent adult neural progenitor cells. Proceedings of the National Academy of Sciences. 2004;101(47):16659–64.

73. Li J, Ma J, Meng G, Lin H, Wu S, Wang J, et al. BET bromodomain inhibition promotes neurogenesis while inhibiting gliogenesis in neural progenitor cells. Stem cell research. 2016;17(2):212–21.

74. Le Douarin N, Kalcheim C. The neural crest: Cambridge university press; 1999.

75. Matatall KA, Agapova OA, Onken MD, Worley LA, Bowcock AM, Harbour JW. BAP1 deficiency causes loss of melanocytic cell identity in uveal melanoma. BMC cancer. 2013;13:1–12.

76. Torre D, Lachmann A, Ma’ayan A. BioJupies: automated generation of interactive notebooks for RNA-Seq data analysis in the cloud. Cell systems. 2018;7(5):556–61. e3.

77. Zhou Y, Zhou B, Pache L, Chang M, Khodabakhshi AH, Tanaseichuk O, et al. Metascape provides a biologist-oriented resource for the analysis of systems-level datasets. Nature communications. 2019;10(1):1523.

78. Abcam. Immunocytochemistry Protocol: Abcam; 2022 [Available from: https://www.abcam.com/en-us/technical-resources/protocols/icc-protocol?srsltid=AfmBOoqyt4JzQIg_lE0OOXDD4ho0XaMzPhyNsZIHA97qu47Q0IKgCbGx.

