## Supplementary figures and images for "Identification of targetable epigenetic vulnerabilities for uveal melanoma"

### Supplemental Material- Uncropped Westerns

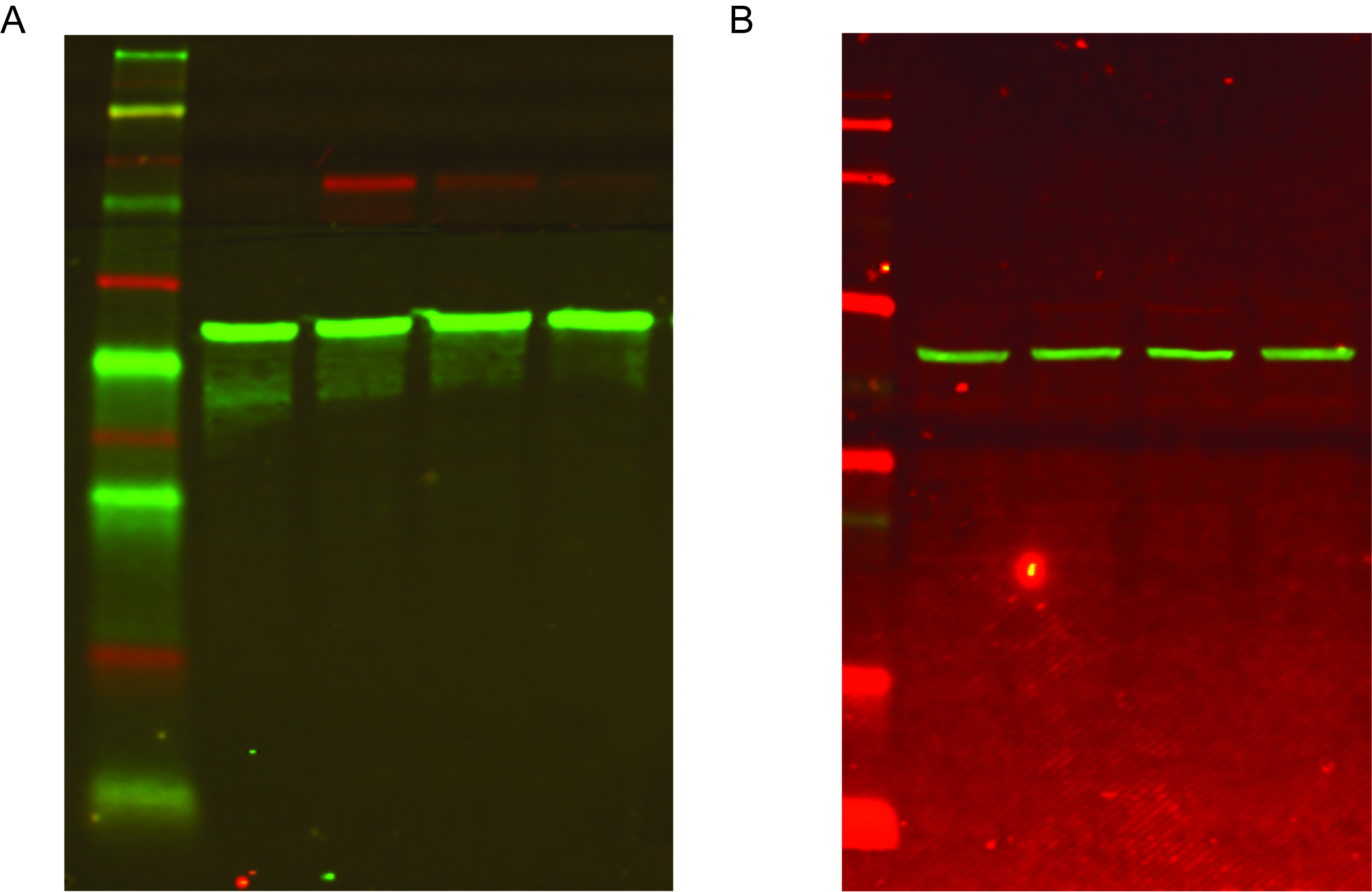
